# DUX4 regulates oocyte to embryo transition in human

**DOI:** 10.1101/732289

**Authors:** Sanna Vuoristo, Christel Hydén-Granskog, Masahito Yoshihara, Shruti Bhagat, Lisa Gawriyski, Eeva-Mari Jouhilahti, Anastassius Damdimopoulos, Vipin Ranga, Mahlet Tamirat, Mikko Huhtala, Kosuke Hashimoto, Kaarel Krjutškov, Gaëlle Recher, Sini Ezer, Priit Paluoja, Pauliina Paloviita, Yujiro Takegami, Ai Kanemaru, Karolina Lundin, Tomi Airenne, Timo Otonkoski, Juha S. Tapanainen, Hideya Kawaji, Yasuhiro Murakawa, Thomas R. Bürglin, Markku Varjosalo, Mark S. Johnson, Timo Tuuri, Shintaro Katayama, Juha Kere

## Abstract

During the human oocyte-to-embryo transition, the fertilized oocyte undergoes final maturation and the embryo genome is gradually activated during the first three cell divisions. How this transition is coordinated in humans is largely unknown. We show that the double homeodomain transcription factor DUX4 contributes to this transition. DUX4 knockdown in human zygotes caused insufficient transcriptome reprogramming as observed three days after fertilization. Induced DUX4 expression in human embryonic stem cells activated transcription of thousands of newly identified bi-directional transcripts, including putative enhancers for embryonic genome activation genes such as LEUTX. DUX4 protein interacted with transcriptional modifiers that are known to couple enhancers and promoters. Taken together, our results reveal that DUX4 is a pioneer regulating oocyte-to-embryo transition in human through activation of intergenic genome, especially enhancers, and hence setting the stage for early human embryo development.

---

Mammalian pre-implantation development commences with oocyte-to-embryo transition, which involves fundamental changes in the epigenetic landscapes, modulation of cell cycle control, and translation or clearance of selected maternal mRNAs, culminating to embryonic genome activation^1,2^. The pioneer regulators orchestrating the oocyte-to-embryo transition and first embryo genome activation steps in human remain poorly understood. The conserved double homeodomain transcription factor *DUX4* represents a plausible candidate regulating the oocyte-to-embryo transition in humans, given its capacity to activate germline genes and genomic repeat elements^3–5^. Here we show that *DUX4* is able to launch the first reprogramming steps from oocyte to embryo in human by activating thousands of novel enhancers and therein, modulating the transcriptome and chromatin. Human *DUX4* knockdown embryos are viable until the third day of development, but their transcriptome is severely altered. Our proteomics approaches suggest that DUX4 binds to Mediator complex and chromatin modifiers through its C-terminal domains, providing a likely explanation to how DUX4 may extensively modulate the genome. This study implies a wider role for *DUX4* as a cellular gate keeper acting both as a general genomic modifier of cell fate as well as a specific inducer of first wave embryo genome activation genes.

## Results

### Quantification of DUX4 in human embryos

The DUX4 induced gene network is highly conserved^6^ and recent reports showed that DUX4 is expressed in early human embryos^3,4^. However, details of this dynamic process, including initiation of *DUX4* expression, remained ambiguous. Therefore, we first set out to quantify *DUX4* mRNA expression levels in human metaphase II (MII) oocytes, zygotes, and cleavage embryos as well as DUX4 protein levels in human zygotes and early embryos (Fig. 1a). We found significant *DUX4* mRNA upregulation in zygotes, while few transcripts were found in MII oocytes or cleavage embryos^7^ (Fig. 1b). Induction of the *DUX4* mRNA orthologues in mouse and non-human primate zygotes is evolutionary conserved (Extended Data Fig.1a). Antibody staining revealed that DUX4 protein is highly abundant in the cytoplasm and nuclei of zygotes as well as 2-cell and 4-cell embryos (Fig. 1c). Quantification of the nuclear DUX4 staining intensities in 3D showed a variable but increasing nuclear signal from the zygotes up to 4-cell embryos, while almost no signal was detected in 8-cell embryos (Fig 1c, d). In one single very early 8-cell stage embryo there was high variability in the nuclear DUX4 staining, consistent with rapid clearance of the DUX4 protein (Extended Data Fig. 1b). These results demonstrate that *DUX4* transcripts appear around the time of fertilisation and is followed by cytoplasmic and nuclear localisation of the DUX4 protein during the first two days of human embryo development coinciding with the timeframe of the oocyte-to-embryo transition and activation of the genome.

**Figure 1.**
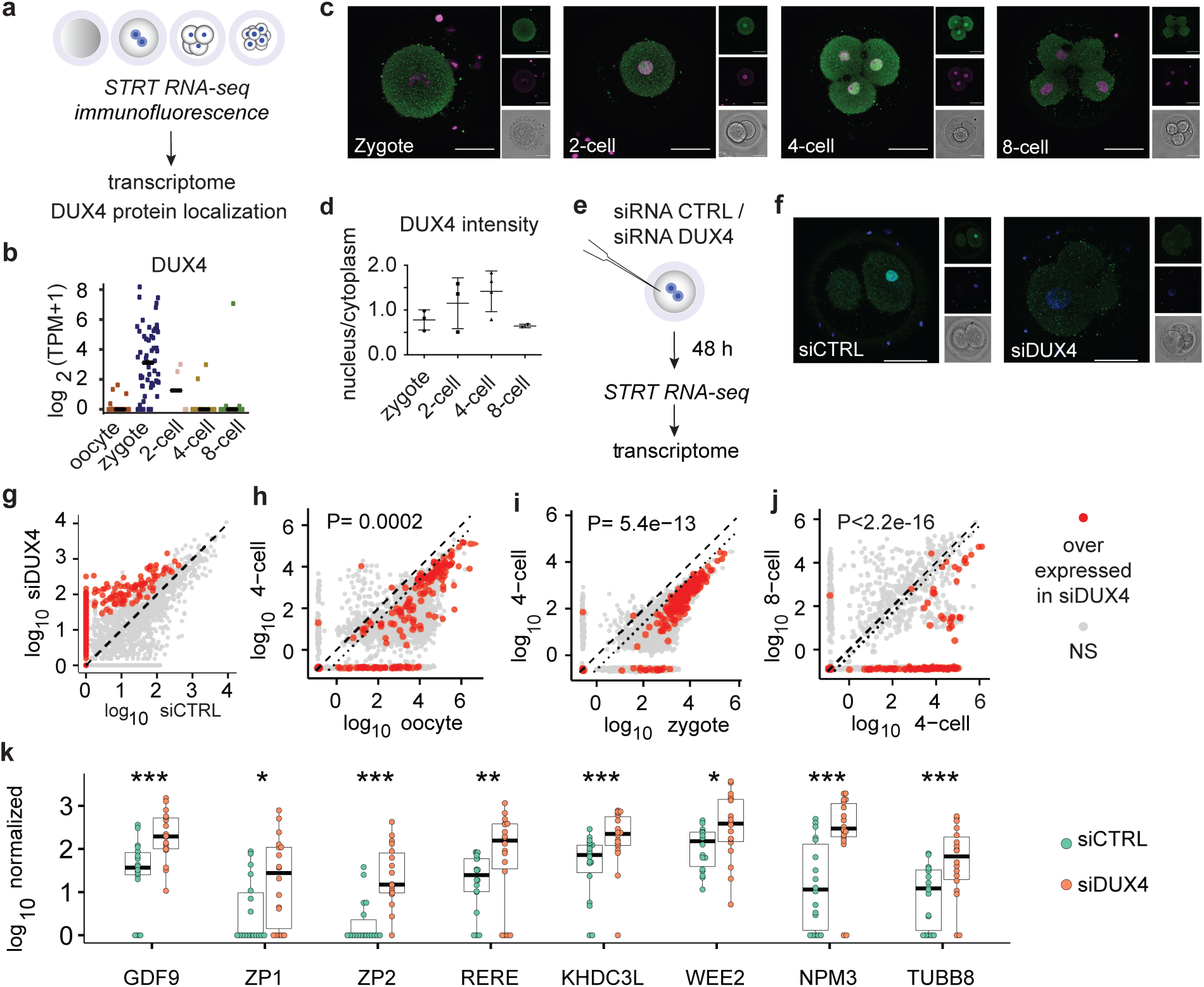
DUX4 knockdown leads to dysregulation of the maternal transcriptome in the human embryo. (a) Schematic of the experimental design. (b) Log2 TPM (Transcripts Per Kilobase Million) of DUX4 mRNA reads in human MII oocytes (n=20), zygotes (n=59), 2-cell (n=4), 4-cell (n=15), and 8-cell (n=14) embryos8. (c) Representative confocal images of zygotes (n=3), 2-cell (n=3) 4-cell (n=4), and 8-cell (n=2) human embryos stained with monoclonal DUX4 antibody E5-5 (green). Nuclei counterstained with DAPI (magenta). The larger left image of each panel shows the composite of the two small fluorescent images on the right for each stage. (d) Quantification of the DUX4 staining intensity in the nucleus normalized to the intensity in the cytoplasm. The samples as in 1c. Data are mean ±SD. (e) siRNA experimental design. (f) Representative images of human embryos immunostained with DUX4 antibody (green) 24 h after microinjection with either control siRNA (n=4) or DUX4 targeting siRNA (n=5). Nuclei counterstained with DAPI (blue). Left side of each panel is a composite of individual corresponding z planes for DUX4 staining, nuclear staining, and the bright field channel (shown on the right side). Scale bar 50 μm. (g) Scatter plot of the expression levels of TFEs of siCTRL (n= 18 cells from two embryos) versus siDUX4 embryos (n=18 cells from three embryos). Red dots represent significantly upregulated TFEs in siDUX4 embryos while grey dots represent TFEs with no significant change. (h-j) Using the oocyte to embryo transition data from Töhönen et al. 2015, we identified the TFEs present in our siCTRL versus siDUX4 data (g) and plotted them as follows h: oocyte to 4-cell, i: zygote to 4-cell, and j: 4-cell to 8-cell. The dotted line marks the cell division effect on cellular RNA content. The red dots are the upregulated TFEs identified in g. Note that they are downregulated in 4 cell embryos versus oocyte or zygote, and 8 cell versus 4 cell embryos, while in the DUX4 knock-down (siDUX4, g) they remain high, i.e. are not down regulated. P-values were calculated with Fisher’s exact test for the frequency of the siDUX4-upregulated TFEs of the TFEs normally downregulated during respective stages. (k) Expression levels of selected oocyte-specific genes in siCTRL and siDUX4 embryos. Wilcoxon test. Asterisks represent statistical significant changes. ***q<0.001; **q<0.01; *q<0.05. Horizontal lines represent the median values in each group.

### Inhibition of DUX4 in human zygotes

Given the short-term and precise manifestation of *DUX4* mRNA and protein in human zygotes and early cleavage stage embryos, we next asked how *DUX4* regulates the first steps of human embryo development. We microinjected either *DUX4* targeting siRNA (siDUX4) or control siRNA (siCTRL) into triploid human zygotes and followed their development for 48 h after the microinjections, until the third day of development (Fig. 1e). Staining of the DUX4 protein was very faint or absent in the siDUX4 embryos but strongly positive in the siCTRL embryos 24 h after microinjection (Fig. 1f), confirming that the *DUX4* targeting siRNA efficiently down-regulated *DUX4* expression. The cells from the microinjected embryos were collected 48 h after microinjections and sequenced for identification of transcript far 5’-ends (TFEs), which represent transcription start sites of polyA-tailed RNAs^8^. Group-wise comparison suggested that a number of TFEs were upregulated in the siDUX4 embryos (Fig. 1g, Supplementary Information 1). We annotated the upregulated TFEs and compared them to our published gene expression data set^7,8^ on human MII oocytes, zygotes, and cleavage cells. These analyses revealed that a large number of mRNAs enriched in siDUX4 embryos were normally down-regulated during transition from oocytes or zygotes to 4-cell embryos (Fig. 1h, i) and from 4-cell embryos to 8-cell embryos (Fig 1j). The most differentially expressed genes between the siCTRL and siDUX4 blastomeres were maternal genes, such as *GDF9*^9^ and *ZP1* and *ZP2*^10^ (Fig. 1k). In agreement with the presence of maternal transcripts, gene expression enrichment analysis using TopAnat^11^ for the genes retained in the siDUX4 embryos resulted in terms such as ‘female germ cell’ and ‘oocyte’ (Extended Data Fig. 2a). Thus, maternal genes that normally undergo targeted clearance during the oocyte-to-embryo transition were retained after the knock-down of DUX4 in human zygotes. Investigating further, we identified 3,196 TFEs that are highly variable between the microinjected blastomeres. Weighted correlation network analysis (WGCNA)^12^ classified these variable TFEs into three modules: TFEs in blue and brown modules overlapped with the upregulated TFEs during normal pre-implantation development^8^, suggesting embryonic genome activation modules, while the TFEs in the turquoise module were associated with maternal genes (Extended Data Fig. 2b). According to the representative expression pattern, siDUX4 blastomeres did not upregulate the genome activation module TFEs (Extended Data Fig. 2c), suggesting insufficient activation of their genome due to *DUX4* knock-down. Although retroelement-derived transcription by DUX4 binding has been reported^13^, only a few genome activation module TFEs overlapped with the 56 families of repeat elements and known DUX4 binding sites (Extended Data Fig. 2d). Surprisingly, TFEs in the maternal turquoise module overlapped with ERVL/ERVL-MaLR elements and the known DUX4 binding sites (Extended Data Fig. 3d; *P*<0.05 by Fisher’s exact test, the odds ratio >1; *P*-values were corrected by Benjamini-Hochberg procedure). However, they only represented up to 5% of the maternal TFEs. Therefore, DUX4 does not seem to regulate maternal TFEs (turquoise module) through ERVL-MaLR promoter elements.

### Transcriptome changes induced by DUX4

To investigate *DUX4* functions, we transfected two human embryonic stem cell (hESC) lines H1 and H9 with DUX4-EmGFP TetOn constructs (Fig. 2a and Extended Data Fig. 3a-d) and analysed transcriptome and chromatin status after inducing *DUX4* expression using doxicycline. Of the previously reported 32 minor genome activation TFEs^8^, 23 (~72%) including *ZSCAN4, TRIM48, LEUTX*^14^, and 3 previously unannotated genes (Extended Data Fig. 4) were significantly upregulated in the EmGFP (+) cells (Fig. 2b, Supplementary Information 2). About 74% (17/23) of the promoters of these TFEs contained *DUX4* binding sites^5,13^ (Fig. 2c). Both, a *de novo* DNA motif, which was highly similar to the known *DUX4* motif, and the known binding site were enriched at the proximal upstream sequence of the upregulated TFEs (Fig. 2d, e). Furthermore, the promoter regions of these transcripts were remarkably overrepresented with DUX4 binding sites among hundreds of transcription factors (Extended Data Fig. 3e). In contrast, only ~11% (14/128) of the major embryo genome activation TFEs were up-regulated (Fig. 2c), suggesting that DUX4 acts only as an inducer of the minor genome activation genes. According to the classification of the differentially regulated TFEs, the vast majority of the upregulated TFEs were mapped to intergenic regions of the genome and the majority of the downregulated TFEs were mapped to the coding regions of the genome (Fig. 2f). We compared these unannotated TFEs upregulated by DUX4 induction with FANTOM-CAT non-coding RNA database and found that 430 TFEs overlapped with 394 long non-coding RNA (lncRNA) exons out of which 46 were antisense lncRNAs. About 42% (1,844/4,415) of the TFEs overlapped with the ERVL-MaLR elements. Finally, we studied the chromatin status of the EmGFP (+) and EmGFP (-) *DUX4* TetOn hESCs using ATAC-sequencing. *DUX4* caused rapid chromatin opening (hereafter referred to as ATAC-gained) in the EmGFP (+) cells. Out of these ATAC-gained chromatin sites, 48.9% significantly overlapped with ERVL-MaLR elements and they were enriched for the DUX4 binding sites (55.8% *P* < 2.2e-16) (Fig. 2g). The ATAC-gained ERVL-MaLR regions remarkably overlapped with the open chromatin regions found in 2-cell human embryos^15^ (Extended Data Fig. 3f). Out of the DUX4-induced gained chromatin regions that overlapped with those of the 2-cell embryos and DUX4 binding sites, 76.7% were unannotated. This suggests that DUX4 is a strong modulator of the intergenic genome immediately after fertilization.

**Figure 2.**
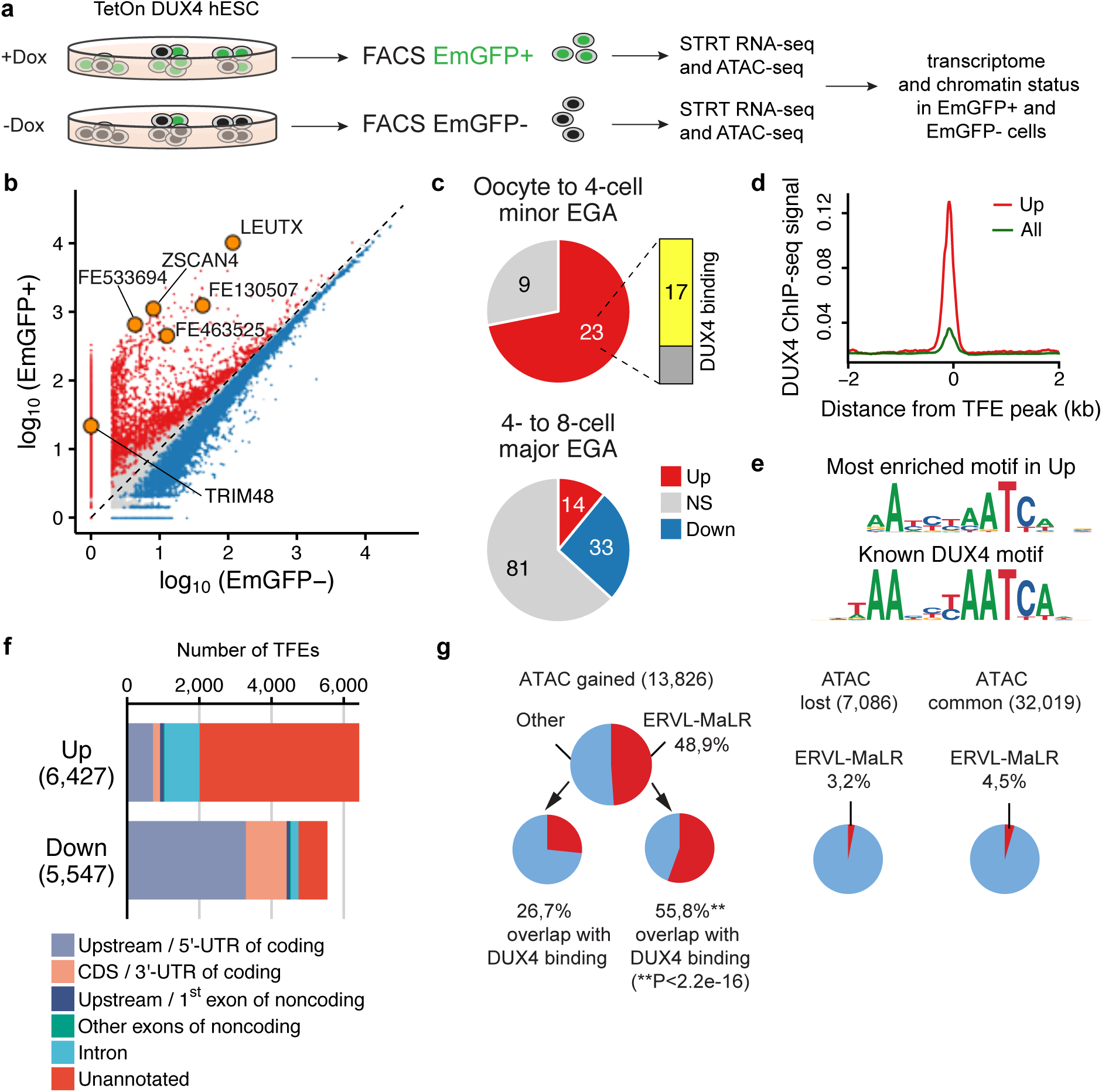
DUX4 causes upregulation of intergenic genome regions and minor embryo genome activation transcripts in human embryonic stem cells. (a) Schematic overview of the experimental design. (b) Overexpressed (red), downregulated (blue), and non-significantly changed (grey) TFEs after DUX4 induction (EmGFP (+) cells. Three samples from two clones of each DUX4 TetOn hESC lines (H1 and H9) were FACS sorted and collected per indicated condition. (c) Proportion of the upregulated (Up), downregulated (Down), and non-significantly changed (NS) TFEs upon DUX4 induction as in (b) among the minor (oocyte to 4-cell embryo) and major (4- to 8-cell embryo) embryonic genome activation genes. One TFE out of the 129 major EGA genes annotated on an unassigned chromosome (ChrUn) and was excluded from the analysis. (d) DUX4 ChIP-seq intensity5 around the peaks of reads within the upregulated TFEs (red) and all the detected TFEs (grey). (e) *De novo* motif enrichment analysis of the DUX4-induced TFEs. Upper panel: the most significantly enriched motif (*P* = 1e-961) in upregulated (UP) genes using binomial statistical test. Lower panel: the best-matching known binding motif of DUX4 (DUX4 ChIP-seq of myoblasts: GSE7579179; matching score = 0.92). (f) Positional information of the upregulated and downregulated TFEs after DUX4 induction. (g) Proportion of the gained, lost, and common ATAC-seq peaks overlapping ERVL-MaLR regions after DUX4 induction. The samples as in b.

### Induction of embryonic genome activation by DUX4 driven enhancers

Substantial upregulation of intergenic genomic regions after *DUX4* induction (Fig. 2f) prompted us to study transcribed enhancers in the *DUX4* TetOn hESCs (Fig. 3a). For this, we investigated native elongating transcripts using cap analysis of gene expression (NET-CAGE)^16^, which sensitively identifies unstable transcripts such as enhancer RNAs (Extended Data Fig. 5a, b). Integration of our TetOn *DUX4* hESC ATAC-seq and NET-CAGE datasets (Fig. 2a and 3a) showed that open chromatin regions were highly enriched at the nucleosome depleted regions of *DUX4* induced (Dox+) enhancers but not at enhancers identified in the Dox (-) hESCs (Extended Data Fig. 5c). Altogether, we identified more than 10,000 transcribed enhancers in our Dox (+) *DUX4* TetOn hESCs and ~ 90% of these enhancers have not been identified previously^16–18^ (Fig. 3b, Supplementary Information 3). Hereafter, novel enhancers that are exclusive to Dox (+) *DUX4* TetOn hESCs are called “novel DUX4 enhancers”. Notable, 36.7% of the novel *DUX4* enhancers overlapped with ERVL-MaLR elements (Fig. 3b). We next annotated enhancer expression in our data and identified putative enhancers for three upregulated minor genome activation genes; *LEUTX* (Fig. 3c, d), previously unannotated *RETT FINGER PROTEIN* (Fig. 3c, Extended Data Fig. 4), and either for *KHDC1* or *KHDC1L* (Fig. 3c, Extended Data Fig. 4). Further, promoters of 12 minor embryo genome activation genes were significantly upregulated by *DUX4* induction (Fig. 3c, Supplementary Information 4). We identified the exact promoter position for the *ZSCAN4* that is upregulated by *DUX4* expression (Extended Data Fig. 5d). We designed guide RNAs for the *LEUTX* promoter and putative novel enhancer regions (Extended Data Fig. 5e) to experimentally test these using the CRISPRa activation system^19^ in HEK293 cells. The expression level of *LEUTX* nearly doubled when the guide RNAs targeting the promoter region were transfected together with the putative enhancer 1 targeting guide RNAs in comparison to the promoter targeting guide RNAs only (Fig. 3e). Out of the 56 retroelement families ERVL-MaLRs significantly overlapped with the novel DUX4 enhancers (Fig. 3f and Extended Data Fig. 5f), constituting ~37% of all novel DUX4 enhancers (*P*<2.2e-16; Fig. 3b). Out of novel DUX4 enhancers 28% overlapped with DUX4 binding sites (*P*<2.2e-16). Using DUX4 ChIP-seq data we compared whether ERVL-MaLRs regions were more often associated with gene promoter or enhancer regions. Only 9.5% of the DUX4 binding site overlapping ERVL-MaLRs were associated with promoter regions while ~37% were associated with enhancer regions (Fig. 3g). In summary, *DUX4* induces a large number of novel enhancers, many of which overlap with ERVL-MaLR regions and regulate the genome at the time of oocyte-to-embryo transition.

**Figure 3.**
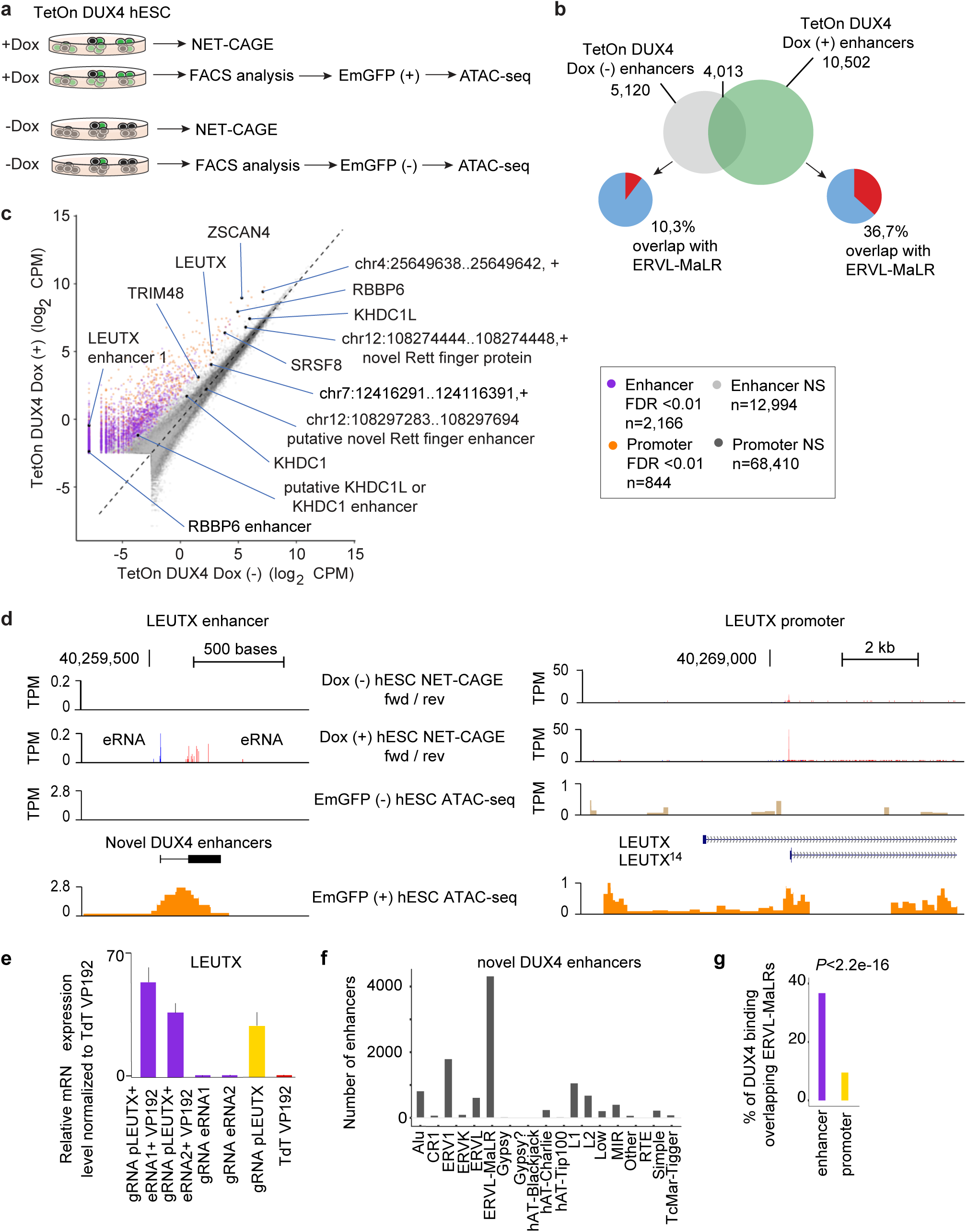
DUX4 activates newly identified enhancers of the minor embryo genome activation transcripts. (a) Experimental design. (b) Venn diagrams of the novel transcribed enhancers identified in TetOn DUX4 Dox (-) (n=two biologically independent samples) and TetOn DUX4 Dox (+) (n=two biologically independent samples) cells. Pie charts show percentage of enhancers overlapping ERVL-MaLR repeat regions. (c) Comparison of TetOn DUX4 Dox (-) hESCs and TetOn DUX4 Dox (+) hESCs NET-CAGE data for promoters and enhancers. Lowly expressed promoters and enhancers with average expression < −2.5 log2 CPM in TetOn DUX4 Dox (-) hESCs (n=2) and TetOn DUX4 Dox (+) hESCs (n=2) were filtered out. Yellow dots, differentially transcribed promoters (FDR ≤ 0.01); purple dots, differentially transcribed enhancers (FDR ≤ 0.01); dark grey dots, non-significant promoters; light grey dots, non-significant enhancers; black dots, promoters and enhancers for minor embryo genome activation genes. (d) NET-CAGE data shows bidirectional transcription for the putative LEUTX enhancer after DUX4 induction (the Dox (+) cells). ATAC-seq data illustrates open chromatin at the putative LEUTX enhancer region after DUX4 induction (the EmGFP (+) cells). The LEUTX promoter was activated after DUX4 induction (the Dox (+) cells, correlating with open chromatin region (ATAC-seq peaks in the EmGFP (+) cells). (e) Relative expression level of LEUTX in HEK293 cells transfected with the indicated guide RNAs and plasmids. gRNA, guide RNA; eRNA, enhancer RNA; VP192; dCas9 transactivator plasmid; pLEUTX, LEUTX promoter. (f) Novel DUX4 enhancers overlapping with retroelement families. The retroelement families overlapping with at least ten DUX4 enhancers are shown. (g) Proportion of the DUX4 binding sites overlapping ERVL-MaLRs at enhancer or promoter regions.

### DUX4 protein domains mediating the DUX4 interactions

Given predominant DUX4 protein presence in the embryos and stage-specific nuclear localization, we set out to study how *DUX4* could mediate such a powerful induction of the genome. For this, we analysed the structural features and protein-protein interactions of DUX4. DUX4 comprises two homeodomains and an intrinsically disordered region with three regions of predicted low disorder conserved in primates. Two predicted amphipathic helices contain a nine amino acid transactivation domain (9aaTAD^20^), also present in LEUTX^21^, and a motif known to recruit the KIX domain^22^ of the cAMP-response element binding protein (CREB)-binding protein (CBP)^23^ (Fig. 4a). We modelled the 9aaTAD peptide 371GLLLDELLA379 and the 416EYRALL421 peptide (KBM, KIX binding motif) into the MLL and pKID/c-Myb site of the ternary complex NMR structure of human KIX from CBP^24^ (PDB: 2LXT) (Fig. 4b, Supplementary Information 6). The hydrophobic residues of 9aaTAD and KBM complement well what is seen in the KIX:MLL:pKID complex. Indeed, experimental tight binding (Extended Data Fig. 6 a-c) was detected for peptides overlapping the 9aaTAD (K_d_ ≈ 0.2 µM) and KBM (K_d_ ≈ 0.6 µM) sequences of DUX4 to KIX domain, and for KBM binding in the presence of 9aaTAD (K_d_ ≈ 1.1 µM). Because DUX4 is observed in the cytoplasm (Fig. 1c), we asked whether the homeodomain1-linker-homeodomain2 structure would be stabile as a unit without bound DNA and subjected the crystal structure of DUX4 (PDB: 6E8C^25^, Supplementary Information 7) to molecular dynamics simulations. Ten residues, highly conserved in primates, form two interacting clusters (Extended Data Fig. 6 d, e), stabilizing both domains even in the absence of DNA (Supplementary movie 8). While the predominantly charge-charge interactions hold the two homeodomains together (Extended data Fig. 6 f-i), the intermediate linker loop imparts flexibility, which could be vital to accommodate DNA once DUX4 enters the nucleus and locates its binding motif. Indeed, the double homeodomain without DNA opened dramatically, by over 38 Å, and the stabile open conformation would be suited to initial interactions with DNA and be consistent with the proposed two-step clamp-like binding mechanism^26^.

**Figure 4.**
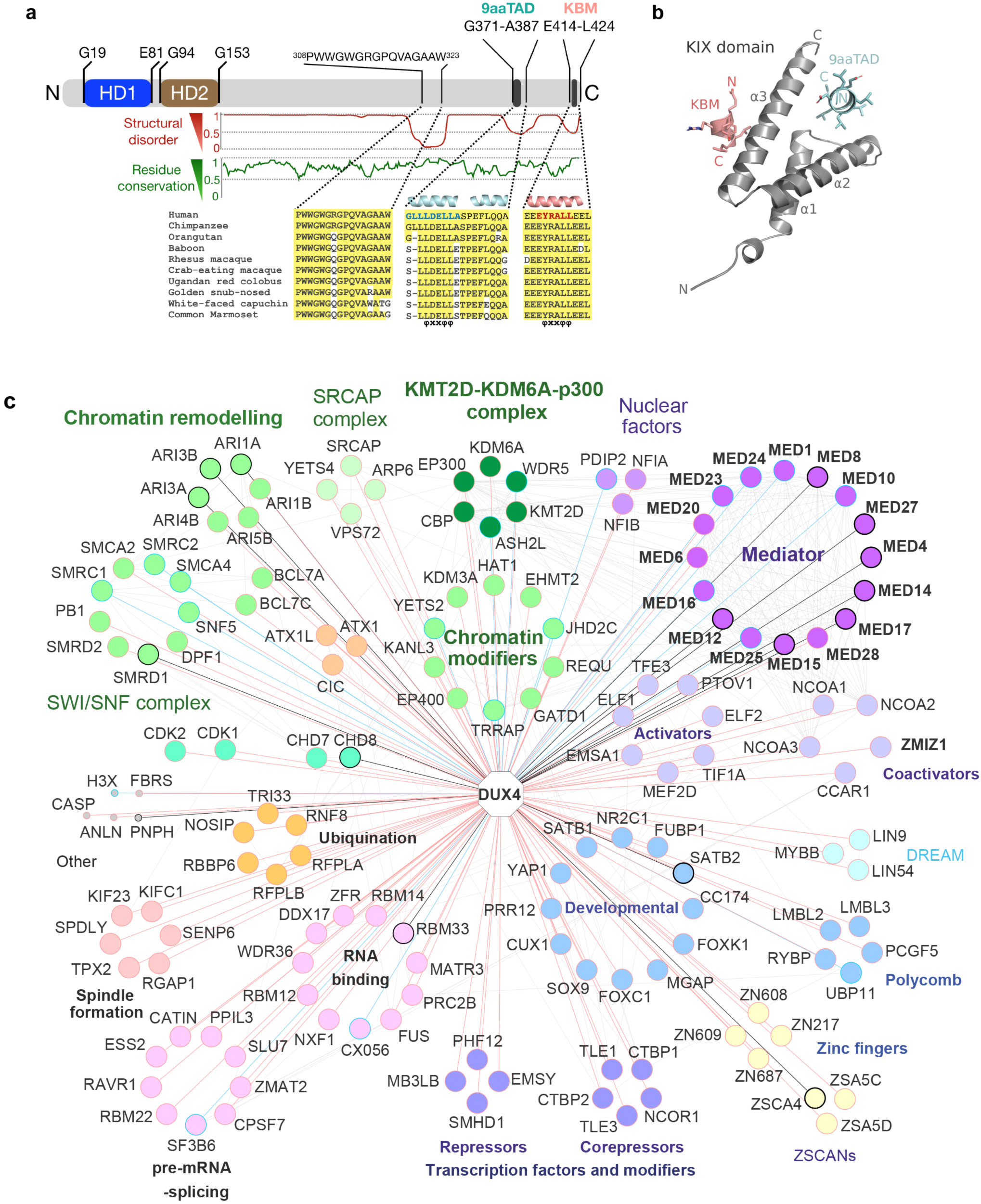
C-terminus of human DUX4 interacts with the transcriptional Mediator complex and chromatin regulators. (a) Domain structure of full-length DUX4: N-terminal homeodomains HD1 and HD2, and C-terminal region. Conservation of residues in primates versus human sequences (green curve) C-terminal to residue G153 and sequence alignment of three conserved regions with a disorder value lower than 0.5 (red curve). Residue numbering from UniProt ID Q9UBX2. Two helical regions are predicted within the C-terminal region, the first one (cyan helices) and the second one (salmon helix) both containing the amphipathic “ΦXXΦΦ” motif (Φ, bulky hydrophobic amino acid; X, any amino acid) found in several transcription factors reported to interact with KIX80-82. The position of the 9aaTAD (blue letters) and KBM (KIX binding motif; red letters) sequences are indicated by black bars. (b) Modelled interactions of the human KIX domain (PDB: 2LXT) with DUX4 9aaTAD (cyan) and KBM (salmon). (c) DUX4 protein-protein interactome. BioID -interactions shown with red lines and AP-MS -interactions with blue lines. If protein appeared in both data sets it is outlined in bold black. Known prey-prey interactions shown in grey (iREF).

To identify protein-protein interactions of DUX4 we used the MAC-tag method where stable (AP-MS) and transient (BioID) interactions can be reliably identified^27,28^. We identified 43 AP-MS and 158 BioID high-confidence DUX4 interactions, out of which 19 appeared in both datasets (FDR < 0.05, corresponding to > 0.73 SAINT Score) (Supplementary Information 9). Overrepresentation Enrichment Analysis (ORA) of protein pathway markers (Reactome, KEGG) showed significant enrichment (*P* < 0.05, FDR < 0.01) of markers linked to ‘transcription’, ‘RNA polymerase II Transcription’, ‘chromatin organization’ and ‘chromatin modifying enzymes’. Comparison of our list of genes to the protein complex databases ComplexPortal and Corum using Fisher’s Exact Test yielded significant overrepresentation of the SWI/SNF chromatin remodelling complex, NSL and NuA4 histone acetyltransferase complex, SRCAP histone exchanging complex, and the Core Mediator complex, (*P* < 0.05, FDR <0.01) (Fig. 4c). Several DUX4 interacting proteins were classified as RNA binding (GO:0003723), spliceosome and pre-mRNA-splicing (Fig. 4c and SI 10). As the protein interaction assay was performed in the HEK 293 cell line, we studied which of the identified DUX4 interacting proteins are expressed by human oocytes or embryos^8,29^. Nearly all genes coding for the DUX4 interacting proteins were expressed in oocytes, embryos, or both. These results suggested that DUX4 may regulate maternal and embryonic proteins in the cytoplasm and nucleus during the oocyte-to-embryo transition and minor embryonic genome activation.

## Discussion

The oocyte-to-embryo transition gradually sets the stage for embryo development^30–33^. DUX4 is an obvious primary candidate mediating chromatin and transcriptome changes that are crucial for oocyte-to-embryo transition. Knock-down of *DUX4* in the human zygotes did not cause mitotic arrest during the 2-day experiment, in agreement with recent findings on *Dux* in mouse embryos where a few of the embryos may proceed until the blastocyst stage^3,34,35^. In the mouse *Dux*-/- embryos, around one third of the embryo genome activation transcripts that are normally upregulated were instead downregulated^35^, while in our human *DUX4* knock-down embryos, many of the maternal, normally downregulated genes remained unchanged. This shows that the role of *DUX4* in the human is not limited to embryo genome activation but that *DUX4* alone is not sufficient for either the oocyte-to-embryo transition or the embryonic genome activation. Given that DUX4 seems to contribute to the maternally biased expressed genes^36^, knockdown at the human zygote stage may hamper observing the entire spectrum of DUX4 functions in the human oocyte-to-embryo transition.

Transient upregulation of *DUX4* mRNA in zygotes and the increasing nuclear DUX4 protein intensity from zygotes until 4-cell stage embryos and its clearance from the nuclei of 8-cell embryos suggested that DUX4 is not only an inducer of the minor embryo genome activation transcripts but that it could also modulate the genome more pervasively and already before genome activation takes place. Ectopic expression of *DUX4* in the hESCs caused extensive chromatin opening, largely at newly identified enhancers and ERVL-MaLR elements. Thirty-seven percent of the novel DUX4 enhancers were suggested to be derived from ERVL-MaLR elements, indicating that some of the ERVL-MaLRs may function as cells stage specific enhancers in human embryos. Indeed, retroelement regions have been suggested to function as regulatory elements, providing novel promoters and possibly enhancers to increase transcriptional complexity especially during development^37,38^. Accordingly, long terminal repeat elements have been suggested as key elements contributing to the oocyte-to-embryo transition^39^. DUX4 upregulation around the time of fertilization may contribute to switching from the maternal^40^ to the cleavage embryo specific retroelements. It is intriguing to speculate, whether activation of ERVL-MaLR elements together with the DUX4 enhancerome provides a correct ‘genomic niche’ for the subsequent genome activation step. It was recently shown that in mouse Dux binding at *Mervl* loci drives chromatin reorganisation at these loci in 2-cell embryo-like cell lines, and that chromatin organisation during early mouse development is a consequence of the *Mervl* integration^41^. To date, human 2-cell-like cell lines have not been established, but importantly, in our experiments, activation of the DUX4 and its likely binding at ERVL-MaLR elements^6^ could modify chromatin towards human cleavage embryo-like stage in the hESCs.

According to our DUX4 proteome data some of the strongest DUX4 interactors were the Mediator complex proteins, many of them identified in both stable and transient interactions. Mediator complex proteins interact with both chromatin modifiers and sequence-specific transcription factors^42^ and they have been shown to connect enhancers to promoters^43–45^. The 9 amino acid transactivator domain present in DUX4 is known to interact specifically with MED15 transcriptional mediator^22^, also present in our proteome interaction data. Our modelling of the DUX4 protein structure suggested that DUX4 C-terminus contains two transcriptional transactivator domains, the 9aaTAD and the KIX-binding domain. According to our protein model, the KIX binding motif of the DUX4 appears highly functional, alluding to DUX4 having all the attributes for rapid target binding and activation, as an ideal candidate for rapid modification of a number of genomic regions, including the newly identified enhancers. Therefore, our data suggests that by using its C-terminal domains, DUX4 binds Mediator complex proteins and chromatin modifiers, like p300^23^, and modulates the transcriptome, including enhancers, during oocyte-to-embryo transition. In conclusion, DUX4 is a pioneering factor participating in setting the stage for human embryo development.

## Online methods

### Human pre-implantation embryos for single cell RNA-sequencing using single-Cell Tagged Reverse Transcription (STRT)

We analysed single cell RNA-sequencing data from Töhönen et al.^8^ for MII oocytes (n=20), zygotes (n=59), 2-cell (n=4), 4-cell (n=15) and 8-cell (n=14) embryos. For the DUX4 knockdown experiment, siCTRL cells (n=18 cells, from two embryos) and siDUX4 cells (n=18 cells, from three embryos) were analysed. The embryos were incubated in Ca^2+^/Mg^2+^-free culture medium (Biopsy Medium, Origio) at 37°C on a heated stage for separation of the cells. Individual cells were briefly rinsed in Ca^2+^/Mg^2+^-free PBS and placed directly in lysis buffer (5mM Tris-HCl, pH 7.0 (LifeTechnologies), 5mM DTT (Thermo Scientific), 0.02% Triton X-100 (Fisher Scientific), 0.5 U/µl Ribolock RNAse inhibitor (Thermo Fisher)). The library was prepared according to the published protocol^8,46,47^. The amplified libraries were sequenced on the Illumina HiSeq2000 instrument.

### Bulk RNA-sequencing of FACS sorted cells using STRT method

TetOn-DUX4 hESCs either with or without doxicycline treatment (see above) were incubated with TrypLE for 5 min, detached, and suspended into cold FACS buffer (5% FBS in PBS). The cell suspension was filtered through Cell strainers to remove any cell clumps and centrifuged at 107*g* pm for 5 min. The cell pellets from Dox (+) and Dox (-) cultures were suspended in the cold FACS buffer and placed on ice. EmGFP (-) cells from the Dox (-) and EmGFP (+) cells from the Dox (+) suspension were sorted into cold FACS buffer using a Sony SH800Z Cell Sorter with blue laser (488) and 100 µm nozzle. Total RNA was isolated from FAC-sorted DUX4-TetOn hES cells using the RNAqueous Total RNA Isolation Kit (AM1912; Thermo Fisher Scientific). 20 ng of total RNA from each sample was used for library preparations. The libraries were prepared using the STRT method as above, with minor modifications. Briefly, RNA samples were placed in a 48-well plate in which a universal primer, template-switching oligos, and a well-specific 8 bp barcode sequence (for sample identification) were added to each well^48,49^. The synthesized cDNAs from the samples were then pooled into one library and amplified by single-primer PCR with the universal primer sequence. The resulting amplified library was then sequenced using the Illumina NextSeq500 instrument.

### Pre-processing of raw STRT RNAseq reads

The sequenced STRT raw reads were processed usin STRTprep^48^ (v3dev branch commit 91a62d2 available at https://github.com/shka/STRTprep/tree/v3dev). The processed nonredundant reads were aligned to the hg19 human reference genome sequences. External RNA Control Consortium (ERCC) spike-in sequences and the human ribosomal DNA unit (GenBank: U13369) with RefSeq transcript alignments as a guide of exon junctions. For gene-based statistics, uniquely mapped reads within (i) the 5’-UTR or the proximal upstream (up to 500 bp) of the RefSeq protein coding genes, and (ii) within the first 50 bp of spike-in sequences, were counted. For TFE-based statistics, the mapped reads were assembled according to the alignments, and uniquely mapped reads within the first exons of the assembled transcripts were counted, as described in Töhönen et al. 2015^8^.

### Downstream STRT RNA-sequencing data analysis

Differentially expressed genes and TFEs between two groups had i) significantly different distributions between the two groups by Wilcoxon statistics with multiple resampling^49,50^ (q-value < 0.05), and ii) significantly larger variation than technical variation, which was estimated by variation of the sequenced spike-in RNA levels^48,51^, among all samples of the two groups (*P*-value < 0.05 adjusted by Benjamini-Hochberg correction). The differential expression was tested by STRTprep pipeline^48^. Enrichment analysis of anatomical terms for the list of upregulated genes by siDUX4 was performed using the TopAnat^49^. All human genes in the Bgee database (https://bgee.org/?page=top_anat)^11^ were used as background. STRT data of the early human embryo were obtained from Töhönen et al. 2015 and 2017^7,8^ and were overlapped with TFEs using the intersectBed function from BEDTools^52^ (v2.27.1). DUX4 ChIP-seq data was obtained from GSE33838^5^ and scores around the FEs were calculated with computeMatrix and visualized with plotProfile from deepTools^53^ (v3.1.3). Motif enrichment was analyzed using the command findMotifsGenome.pl from HOMER^54^ (v4.10.3) with the option “-size -300,100”. Enrichment analysis with publicly available ChIP-seq datasets was conducted with ChIP-Atlas^55^ (http://chip-atlas.org). A total of 7,216 human transcription factor ChIP-seq datasets which had more than 500 peaks were analyzed. Fold enrichment was calculated as (the number of ChIP-seq peaks overlapping with upregulated TFEs / the number of upregulated TFEs) / (the number of ChIP-seq peaks overlapping with all TFEs / the number of all TFEs). *P*-values were calculated with Fisher’s exact test and *Q*-values were calculated with the Benjamini & Hochberg method. After excluding the TFEs annotated on ribosomal DNA, 6,425 upregulated TFEs were used as foreground and 109,624 all the detected TFEs were used as background both in the motif and ChIP-seq enrichment analysis.

### Library preparation, sequencing and read-alignment for CAGE-based data

Nascent RNA from flash-frozen cells was isolated as described by Hirabayashi et al.^16^ with the following exceptions: (i) 5× DNase I enzyme (Thermo Fisher Scientific) was used to prepare the DNase I solution (50 µl), (ii) the samples were incubated for up to 1 h at 37°C while being pipetted up and down several times every 10 min, and (iii) RNA quality was measured using TapeStation 4200 (Agilent). CAGE-based libraries were generated according to the no-amplification non-tagging CAGE libraries for Illumina next-generation sequencers (nAnT-iCAGE) protocol^56^. All CAGE-based libraries were sequenced in single-read mode on an Illumina NextSeq500 platform. Reads were split by barcode using the MOIRAI^57^ package. Cutadapt v 1.1.8 (http://code.google.com/p/cutadapt/) was used to trim reads to 73 bp, and remove reads below base quality 33 and ‘N’ bases. Reads aligning to ribosomal RNA sequences (GenBank U13369.1) were removed using the rRNAdust script within the MOIRAI package. The resulting reads were aligned to the human genome (hg19) using STAR v 2.5.0a^58^ with Gencode v27lift37 (“comprehensive”)^59^ as the reference gene model. Mapping was performed with the following parameters: --runThreadN 12 -- outSAMtype BAM SortedByCoordinate --out FilterMultimapNmax 1. Following alignment, the technical replicates were merged using the Picard Toolkit v 2.0.1 with the MergeSamFiles program (Broad Institute, Picard Toolkit, 2018. http://broadinstitute.github.io/picard).

### Identification of transcribed promoters and enhancers

Reads mapping to known FANTOM5 promoters^60^ and FANTOM-NET enhancers^16^ were counted and normalized essentially as described in^16^. Decomposition peak identification (https://github.com/hkawaji/dpi1/blob/master/identify_tss_peaks.sh) was used to identify tag clusters with default parameters but without decomposition. Peaks with at least three supporting CAGE tags were retained and used as input to identify bidirectional enhancers (https://github.com/anderssonrobin/enhancers/blob/master/scripts/bidir_enhancers). Differential expression (DE) analysis with Benjamini–Hochberg false discovery rate (FDR) correction was performed for promoters and enhancers using egdeR v3.26.8^61,62^. Lowly expressed promoters and enhancers (average value between replicates < −2.5 log_2_ CPM) were excluded from the analysis. Promoters and enhancers with FDR ≤ 0.01 were identified as differentially expressed.

### LEUTX enhancer validation

Putative LEUTX enhancer regions 1 and 2 were predicted from TetOn DUX4 hESC NET-CAGE dataset. The guide RNAs targeting the LEUTX promoter and each of the putative enhancers were designed using the Benchling CRISPR tool (https://benchling.com), targeting them to the proximal promoter (−400 to −50 base pairs from transcription start site) or the putative enhancers. Possible guide sequences were selected according to their off-target score and position. Guide RNA oligos are shown in Extended Data table 7. Guide RNA transcriptional units (gRNA-PCR) were prepared by PCR amplification and transfected to HEK293 cells as described in Balboa et al. 2015^63^.

### Guide RNA production

Guide RNA transcriptional units (gRNA-PCR) were prepared by PCR amplification with Phusion polymerase (Thermo Fisher), using as template U6 promoter and terminator PCR products amplified from pX335 together with a guide RNA sequence-containing oligo to bridge the gap. PCR reaction contained 50 pmol forward (Fw) and reverse (Rev) primers, 2 pmol guide oligo, 5 ng U6 promoter and 5 ng terminator PCR products in a total reaction volume of 100µl. PCR reaction program was 98C/10sec, 56C/30sec, 72C/12sec for 35 cycles. Amplified gRNA-PCRs were purified. When needed, alternative Fw and Rev primers were used to incorporate suitable restriction sites for gRNAPCR concatenation. LEUTX promoter gRNA-PCR units were concatenated using Golden Gate assembly^64^. Destination vector GGdest-ready was generated by PCR-cloning Esp3I destination cassette from pCAG-T7-TALEN (Sangamo)-Destination (Addgene: 37184^65^ into pGEM-4Z (Promega). Assembly reactions contained 150 ng of GGdest-ready vector, 50 ng of each gRNA-PCR product (five in total), 1 uL Esp3I (Thermo Fisher, ER0451), 2 uL T4 DNA ligase (Thermo Fisher, EL0011), 2 uL T4 ligase buffer and 2 uL DTT (10mM, Promega, V3151) in a final volume of 20 uL. Thermal cycle consisted of 50 cycles of restriction/ligation (2 min at 37°C, 5 min at 16°C) followed by enzyme inactivation step (20 min at 80°C). Ten microliters of the reaction were transformed into DH5alpha chemical competent bacteria and plated on LB agar containing ampicillin. Correct concatenation of the gRNA-PCR products was confirmed by sequencing.

### HEK cell transfection

HEK 293 cells were seeded on tissue culture treated 24 well plates one day prior to transfection (10^5^ cells/well). Cells were transfected using FuGENE HD transfection reagent (Promega) in fibroblast culture medium with 500 ng of dCas9VP192 transactivator encoding plasmid and 200 ng of gRNA-PCR or gRNA-PCR and vector containing LEUTX promoter targeting guides. Cells were cultured for 72 hours post-transfection, after which samples were collected for qRT-PCR.

### Human ESC culture

hESC lines H1 (WA01) and H9 (WA09) were purchased from WiCell. The hESCs were maintained on Geltrex-coated tissue culture dishes in Essential 8 culture medium and passaged every three to five days by incubation with 0.5 mM EDTA (all from Thermo Fisher Scientific).

### Plasmid construction

The full-length DUX4 (NM_001293798.2) was synthesized and cloned between the SalI and BamHI sites of the pB-tight-hMAFA-ires-EmGFP-pA-PGK-Puro vector (a kind gift from Diego Balboa, Biomedicum Stem Cell Centre) at GenScript (Genscript, NJ, USA).

### Generation of TetOn DUX4 human embryonic stem cells

hESCs were incubated with StemPro Accutase (Thermo Fisher Scientific) until the edges of the colonies started to curl up. The Accutase was aspirated and the cells were gently detached in cold 5% FBS (Thermo Fisher Scientific) 1×PBS (Corning) and counted. One million cells were centrifuged at 107*g* for 5 min and the pellet was transferred into 120 µl of R-buffer containing 1 µg of pB-tight-DUX4-ires-EmGFP-pA-PGK-Puro, 0.5 µg of pBASE and 0.5 µg of rtTA-M2-IN plasmids. 100 µl of the cell-plasmid suspension was electroporated with two pulses of 1100V, 20 ms pulse width, using Neon Transfection system (Thermo Fischer Scientific). The electroporated cells were plated on Geltrex-coated dishes in Essential 8 medium with 10 µM ROCK inhibitor Y27632 (Selleckhem). The following day, the medium was exchanged with fresh Essential 8 medium without ROCK inhibitor. The cells were selected with Puromycin at 0.3 µg/ml. The TetOn-DUX4 hESC clones were picked manually on Geltrex-coated 96-well plates, expanded and selected again with Puromycin. Appearance of the EmGFP reporter protein was tested using Doxycycline at concentrations ranging from 0.2 µg/ml to 1.0 µg/ml and detected using an EVOS FL Cell imaging system (Thermo Fisher Scientific). For the experiments presented in this paper, the DUX4 TetOn hESCs have been treated with 1µg/ml of doxycycline for 1, 2, or 3 h (qPCR) or 4 h (STRT-RNA seq, ATAC-seq, NET-CAGE) prior to subsequent analyses.

### cDNA cloning of previously unannotated genes

A cDNA library was prepared from a single human 4-cell embryo according to the protocol by Tang et al.^66^ and used for cloning of putative transcripts. Transcripts were amplified using Phusion High-Fidelity DNA polymerase (New England Biolabs) according to manufacturer’s instructions. Predicted KHDC1 pseudo gene 1, putative RING-finger type E3 ubiquitin ligase, and putative RING-finger domain protein encoding genes were amplified using touchdown PCR: 98°C for 30 s; 24 cycles of 98°C for 10 s, annealing for 30 s, temperature decreasing from 63°C to 56°C, 1°C/3 cycles, 72°C for 30 s; 16 cycles of 98°C for 10 s, 55°C for 30 s, 72°C for 30 s; final extension 72°C for 10 min. All PCR products were cloned into pCR4Blunt-TOPO vector using the Zero Blunt TOPO PCR Cloning kit (Invitrogen) and sequences were verified by Sanger sequencing (Eurofins Genomics). Clone sequences are available from the ENA browser at http://www.ebi.ac.uk/ena/data/view/LR694082-LR694089.

### Bioinformatics analysis and molecular dynamics simulations of the DUX4 protein

The sequences of the human DUX family proteins were obtained from the UniProt database (The UniProt Consortium), whereas DUX4 sequences from other primates were retrieved from the non-redundant database of NCBI using blastp^67^ with human DUX4 (UniProt ID: Q9UBX2) as the query sequence (SI 5). Multiple sequence alignment over the full-length sequences was carried out using MAFFT^68^ with default parameters. Secondary structures, solvent accessibility and disordered regions were predicted using SCRATCH^69^ and RaptorX-Property^70^. The 9aaTAD web server (“Most Stringent Pattern”^71^) was used to predict 9aaTAD motifs. The crystal structure of the DUX4 HD1-linker-HD2 fragment bound to DNA (PDB: 6E8C^25^) was obtained from the Protein Data Bank (PDB; ^72^). PyMOL (version 2.4; Schrödinger LLC) and Bodil^73^ were used to analyze inter-HD interactions. For modeling the binding of the 9aaTAD peptide^371^GLL**L**DE**LL**A^379^ and the KBM ^416^E**Y**RA**LL**^421^ peptide of DUX4 onto the KIX domain, the NMR structure (model 1/20) of human KIX in complex with MLL and pKID peptide^24^ (PDB: 2LXT) was chosen as the template; the sequence ^846^PSD**I**MD**FV**L^854^ of MLL and ^13^S**Y**RK**IL**^138^ of pKID were mutated in PyMOL to match the DUX4 sequences ^371^GLL**L**DE**LL**A^379^ and ^416^E**Y**RA**LL**^421^, respectively, and the coordinates of extra residues of the MLL and pKID peptides were removed; PDB coordinates for KIX in complex with DUX4 9aaTAD and KBM peptides in Supplementary Information 6.

### Expression of human KIX domain from CBP, binding of C-terminal peptides

A synthetic, codon-optimized gene in pET100/TOPO vector (Invitrogen GeneArt Gene Synthesis, Thermo Scientific) was used to express the human KIX domain of CBP (residues 587-673; Uniprot Q92793) in *E. coli* BL21 DE3 cells. The expressed construct (14.5 kDa) contained 36 extra N-terminal residues, including a 6xHis tag, the XpressTM epitope and an enterokinase cleavage site, in addition to the KIX domain (86 residues). Transformed *E. coli* were grown with ampicillin selection in 600 ml of ZYM-5025 autoinduction medium^74^ for 10 h at 37°C. The cells were collected by centrifugation at 3,000×*g* for 20 min and stored at −20°C. The pellets were thawed and suspended in buffer A (50 mM Tris, pH 8.0, 500 mM NaCl) with 20 mM imidazole and lysed by sonication. The supernatant was separated from the cell debris by centrifugation (45,000×*g* for 40 min) and applied to a three-step purification protocol using an ÄKTA Pure 25 chromatography system (GE) with a UV detector. First, a Histrap HP (1 ml; GE) column was used for metal-affinity chromatography: the sample was applied and the column was subsequently washed with 25 column volumes (CV) of buffer A with 20 mM imidazole. KIX was eluted with a linear imidazole gradient from 20 mM to 500 mM in buffer A over 15 CV, and the column was then washed with 5 CV of 500 mM imidazole in buffer A. The KIX containing fractions (ca. 7 ml) were identified by UV absorbance at 280 nm, pooled, then dialyzed (30× volume, two exchanges, CelluSep dialysis membrane, MWCO 6-8K; Membrane Filtration Products, Inc.) against buffer B (25 mM CHES, pH 9.0). Second, anion exchange chromatography was performed with a Resource Q column (1 ml; GE). The cleared (3,200×*g* for 15 min) dialysis pool was applied to the column, the column was washed with 20 CV of buffer B, and eluted with a linear gradient from 0 to 1 M NaCl in buffer B over 15 CV. The KIX containing fractions were pooled (ca. 4 ml), then concentrated with an Amicon Ultra-4 centrifugal filter (MWCO 3K; Merck Millipore) to a volume of 0.5 ml. Third, the concentrated sample was applied to a Superdex 75 10/300 GL size exclusion chromatography column (GE) and eluted with buffer C (25 mM Tris, pH 8.4, 150 mM NaCl) using a flow rate of 0.5 ml/min (0.5 ml fractions). The purity of the sample was analyzed with SDS-PAGE and Coomassie staining, and the concentration was verified by measuring the UV absorbance at 280 nm with NanoDrop One (Thermo Scientific).

Binding assays were performed using a Monolith NT(TM) microscale thermophoresis instrument (Nanotemper Technologies). The His-tagged KIX domain was labeled non-covalently using Monolith NT(TM) His-Tag Labeling Kit RED-tris-NTA (1st generation; Nanotemper Technologies) according to manufacturer’s instructions. Monolith NT.Automated Capillary Chips (Nanotemper Technologies) were used to test binding and to determine the affinity of the 9aaTAD ^(371^GLLLDELLA^379^) and KBM (^416^EYRALL^421^) peptides to KIX; the homedomain of human LEUTX with His-Tag was used as a negative control. Peptides were ordered from GenScript and dissolved in deionized water. The final concentration of KIX in the assay was 20 nM, and the concentration of each peptide in a binding test assay was 5 µM (250× molar excess). The KIX protein and the peptide samples were diluted in PBS-Tween (pH 7.4 0.05% v/v of Tween 20) buffer for the assays.

### Molecular dynamics simulations

Based on the DUX4 structure (PDB: 6E8C,^25^), molecular dynamics (MD) simulations, over all atoms, were used to explore the dynamic states of DUX4: (1) double HD complex with (HD1-HD2 + DNA) and (2) without (HD1-HD2) bound DNA, and (3) HD1 + DNA and (4) HD2 + DNA. Prior to the simulations, hydrogen atoms and missing side-chain atoms for R22 of DUX4 were added using Chimera^75^. MD simulations with the AMBER package (version 18; Case, D.A., 2018. The Amber Project, https://ambermd.org/CiteAmber.php) used the ff14SB (for protein;^76^) and OL15 (for DNA^77^) force fields. The structures were solvated with explicit TIP3P water molecules^78^ within an octahedral box ensuring a 12 Å distance between the boundaries of the simulation box and solute atoms. Sodium counter ions were added to neutralize the system and additional Na^+^/Cl^−^ ions were added to bring the salt concentration to 150 mM. Periodic boundary conditions were implemented, and the particle-mesh Ewald algorithm was applied^79^ for electrostatic interactions with a distance cutoff of 9 Å. For full details of the simulation protocol see Tamirat et al., 2019^80^. Conformations were saved every 10 ps and the resulting MD trajectories were analysed further by calculating the root-mean-square deviations (RMSD; over backbone atoms) and root-mean-square fluctuations (RMSF; over Cα atoms), as well as monitoring hydrogen bond interactions using CPPTRAJ^81^ and VMD^82^. Coordinates (PDB format) of DUX4 HD1-HD2 *sans* DNA after 100 ns simulation in Supplementary Information 7.

### Cloning of DUX4 to MAC-tag Gateway® destination vector

DUX4 was first amplified in a two-step PCR reaction from pB-tight-DUX4-ires-EmGFP-pA-PGK-Puro and cloned into a Gateway compatible entry clone using Gateway BP Clonase II (Invitrogen) according to manufacturer’s instructions. The entry clone was further cloned to Gateway compatible destination vectors containing the C-terminal and N-terminal tags as described^28^. Transfection and selection of the Flp-In™ T-REx™ 293 cells (Invitrogen, Life Technologies, R78007, cultured in manufacturer’s recommended conditions) and affinity purification of the final product was done as previously^28^.

### Liquid Chromatography-Mass Spectrometry

Analysis was performed on a Q-Exactive mass spectrometer with an EASY-nLC 1000 system via an electrospray ionization sprayer (Thermo Fisher Scientific), using Xcalibur version 3.0.63. Peptides were eluted from the sample with a C18 precolumn (Acclaim PepMap 100, 75 µm × 2 cm, 3 µm, 100 Å; Thermo Scientific) and analytical column (Acclaim PepMap RSLC, 65 µm × 15 cm, 2 µm, 100 Å; Thermo Scientific), using a 60 minute buffer gradient ranging from 5% to 35% Buffer B, then a 5 min gradient from 35% to 80% Buffer B and 10 minute gradient from 80% to 100% Buffer B (0.1% formic acid in 98% acetonitrile and 2% HPLC grade water). 4 µl of peptide sample was loaded by a cooled autosampler. Data-dependent FTMS acquisition was in positive ion mode for 80 min. A full scan (200-2000 m/z) was performed with a resolution of 70,000 followed by top10 CID-MS^2^ ion trap scans with a resolution of 17,500. Dynamic exclusion was set for 30 s. Database search was performed with Proteome Discoverer 1.4 (Thermo Scientific) using the SEQUEST search engine on the Reviewed human proteome in UniProtKB/SwissProt databases (http://www.uniprot.org, downloaded Nov. 2018). Trypsin was selected as the cleavage enzyme and maximum of 2 missed cleavages were permitted, precursor mass tolerance at ±15 ppm and fragment mass tolerance at 0.05 Da. Carbamidomethylation of cysteine was defined as a static modification. Oxidation of methionine and biotinylation of lysine and N-termini were set as variable modifications. All reported data were based on high-confidence peptides assigned in Proteome Discoverer (FDR < 0.05).

### Identification of statistical confidence of interactions

Significance Analysis of INTeractome (SAINT^83^)-express version 3.6.3 and Contaminant Repository for Affinity Purification (CRAPome, http://www.crapome.org) were used to discover statistically significant interactions from the AP-MS data^84^. The DUX4 LC-MS data was ran alongside a large dataset of other transcription factors, as well as a large GFP control set. Final results represent proteins with a SAINT score higher than 0.73, and present in all four replicates.

### Overrepresentation Analysis

Overrepresentation analysis of statistically significant interactions in Gene Ontology and Reactome was done in WebGestalt^85^, and overrepresentation of prey proteins in ComplexPortal^86^ (https://www.ebi.ac.uk/complexportal) and CORUM^87^ (https://mips.helmholtz-muenchen.de/corum/) was done using Fisher’s exact test and multiple testing correction in an in-house R-script.

### Interaction network

Protein interaction networks were constructed from filtered SAINT data that was imported to Cytoscape 3.6.0. Known prey-prey interactions were obtained from the iRef database (http://irefindex.org).

### RNA isolation, reverse transcription and quantitative real-time PCR from DUX4 TetOn hESCs

Total RNA was isolated using NucleoSpin RNA kit (Macherey Nagel). 1µg of RNA was reverse transcribed by MMLV-RTase with oligo dT, dNTPs, and Ribolock in MMLV-RTase buffer (Thermo Fisher Scientific). 5× HOT FIREPol qPCR Mix (Solis Biodyne) was used to measure relative mRNA levels with LightCycler (Roche). The Δ Δ CT method was followed to quantify the relative gene expression where *CYCLOPHILIN G* was used as endogenous control. Relative expression of each gene was normalized to the expression without doxicycline treatment. The primer sequences are listed in Extended Data Table 7.

### ATAC-sequencing library preparation and data analysis

The ATAC-sequencing libraries were prepared as in ^88^. 5×10^4^ EmGFP (-) and EmGFP (+) TetOn-hESCs (H1 clone 2, H1 clone 8, H9 clone 3 and H9 clone 4) were centrifuged at 500×*g* for 5 min. The pellets were washed in cold 1× PBS by centrifugation at 500×*g* for 5min. Each cell pellet was lysed in 50 µl of cold lysis buffer (10 mM Tris-HCl, pH 7.4; 10 mM NaCl, 3 mM MgCl_2_, and 0.1% IGEPAL CA-630) and centrifuged at 500×*g* at 4°C for 10 min. The pellet was then resuspended in the transposase reaction mix (2.5 µl of transposase in TD buffer (Nextera DNA library preparation kit, Illumina) and incubated at 37°C for 30 min. The reactions were purified through columns and eluted in 20 µl. After addition of the barcode oligos the DNA samples were amplified for 12 cycles (98°C for 10 s, 63°C for 30 s and 72°C for 60 s) in Phusion PCR master mix (Thermo Fisher Scientific). The PCR products were purified through the columns and eluted in 20 µl.

### ATAC-seq data analysis

Bcl files were converted and demultiplexed to fastq using the bcl2fastq program. STAR^58^ was used to index the human reference genome (hg19), obtained from UCSC, and align the resulting fastq files. The resulting bam files with the mapped reads were then converted to tag directories with subsequent peaks calling using the HOMER suit of programs^54^. HOMER was also employed for counting the reads in the identified peak regions. The raw tag counts from the peaks were then imported to R/Bioconductor and differential peak analysis was performed using the edgeR package and its general linear models pipeline. Peaks with an FDR adjusted p value under 0.05 were termed significant. Plotting was done in R using packages Complex heatmap, ggplot2 and ggbeeswarm. RepeatMasker table downloaded from UCSC (http://hgdownload.soe.ucsc.edu/goldenPath/hg19/database/rmsk.txt.gz) was converted to BED format and then intersected with the ATAC-seq peaks using the intersectBed from BEDTools^52^ to determine the peaks overlapped with ERVL-MaLR elements. ATAC-seq data of human early embryo were obtained from GSE101571^15^, and scores around the ATAC-seq peaks were calculated with computeMatrix and visualized with plotHeatmap from deepTools^53^. All the scripts and command line options can be provided upon request.

### Immunocytochemistry of human ESC

Cells were fixed with 3.8% PFA, washed three times, permeabilised in 0.5% (v/v) Triton X-100 in PBS for 7 min, and washed with washing buffer (0.1% (v/v) Tween20 in PBS). The samples were incubated with ProteinBlock (Thermo Fisher Scientific) at room temperature for 10 min to prevent unspecific binding of primary antibody. Primary antibody (rabbit MAb anti-DUX4, clone E5-5, Abcam) was diluted 1:300 in washing buffer and incubated at 4°C overnight. After washings, fluorescence-conjugated secondary antibody (anti rabbit 594, A-21207; Thermo Fisher Scientific) was diluted 1:1000 in washing buffer and incubated at room temperature for 20 min. Nuclei were counterstained with DAPI 1:1000 in washing buffer. The images were captured with an Evos FL Cell Imaging system using 10× and 20× Plan Achromatic objectives.

### Immunocytochemistry of human embryos

For characterization and quantitation of the DUX4 protein zygotes (n=3) and embryos (2-cell, n=3; 4-cell, n=4; 8-cell, n=2 plus one early 8-cell stage embryo shown in Extended Data Fig. 1 were fixed in 3.8 % PFA at room temperature for 15 min, washed three times in washing buffer (as above), and permeabilised in 0.5% Triton X-100 in PBS at room temperature for 15 min. Unspecific primary antibody binding was blocked as above. DUX4 antibody (as above) was incubated at 4°C overnight. The embryos were washed and incubated in the secondary antibody (anti-rabbit 488, A-21206; Thermo Fisher Scientific) diluted 1:500 in washing buffer (as above) at room temperature for 2 h. After washings, nuclei were counterstained with DAPI 1:500 in washing buffer. To confirm that DUX4 targeting siRNA efficiently downregulated *DUX4*, siCTRL (n=4) and siDUX4 (n=5) microinjected zygotes were stained for DUX4 as above.

### Imaging and confocal microscopy image analysis

Human embryos were imaged in washing buffer on Ibidi 8-well µ slides with a Leica TCS SP8 confocal laser scanning microscope (Leica Microsystems, Mannheim, Germany) using Leica HC PL APO CS2 40×/1.10NA and Leica HC PL APO CS2 63×/1.20NA water objectives. Confocal images were processed using Fiji (http://fiji.sc). For the data presented in Fig 1c and d, images were smoothened using a Gaussian filter (radius = 1 pixel kernel). For the quantification of the DUX4 intensity in the nucleus (Fig. 1d), the DAPI channel was denoised using a rolling ball (radius = 100). The images were smoothened in 3D using a Gaussian filter (radius = 2 pixel kernel) and cell nuclei were segmented. The segmented regions were used to measure average pixel intensity per nucleus in each cell in the DUX4 channel. DUX4 intensity in the nucleus was normalized to intensity of the corresponding cytoplasmic DUX4 staining in the single representative plane.

### Culture and microinjection of human embryos

Human triploid zygotes were warmed using a Gems Warming Set (Genea Biomedx) and cultured in G-TL medium (Vitrolife) in 6%O_2_ and 6% CO_2_ at 37°C. 12 µl of either 20 µM scrambled control siRNA (AM4611, Thermo Fisher Scientific) or DUX4-targeting siRNA (cat. 4457308, Thermo Fisher Scientific) diluted in nucleotide-free H_2_O were mixed with total of 500 ng of GAP-GFP mRNA and centrifuged at maximum speed at 4°C for 10 min. The embryos were microinjected using Eppendorf microinjector and placed in G-TL medium in a Geri dish for 3D time-lapse imaging (Geri incubator, Genea Biomedx, Australia).

### Human embryo live imaging

Imaging of the human triploid embryos was initiated immediately after microinjections (Geri incubator). Images were captured in 3D every 15 min until the embryos were removed for fluorescence staining or termination of the experiment.

### Ethical approvals

Collection and experiments on human oocytes and embryos were approved by the Helsinki University Hospital ethical committee, diary numbers 308/13/03/03/2015 and HUS/1069/2016. Human surplus oocytes, zygotes, and embryos were donated by couples that had undergone infertility treatments at the Reproduction Medicine Unit of the Helsinki University Hospital. The donations were done with an informed consent and patients understood that donating oocytes, zygotes, or embryos is voluntary and does not affect their treatment in any way.

## Acknowledgements

We are grateful for all the couples that donated their surplus oocytes, zygotes, or embryos for this project. We thank the IVF nurses at the Reproductive Medicine Unit of the Helsinki University Hospital for recruiting the couples for the oocyte and embryo donation program, Dr. Diego Balboa-Alonso for the pB-ires-EmGFP-pA-PGK-Puro plasmid and Dr. Jere Weltner for insightful discussions. We acknowledge Biomedicum Imaging Unit (Helsinki), Functional Genomics Unit (Helsinki), Biomedicum Flow Cytometry Unit (Helsinki), Bioinformatics and Expression Analysis Core Facility (Stockholm) for skilled technical assistance. We thank Dr. Jukka Lehtonen for scientific IT support (Biocenter Finland) and CSC IT Center for Science for supercomputing. We thank prof. Outi Hovatta for inspiration and introducing the senior author to the field of early embryogenesis. This project was supported by Jane and Aatos Erkko foundation, Sigrid Jusélius foundation, Finnish Academy, and Helsinki University Hospital funds. SV was supported by Jane and Aatos Erkko foundation. MY was supported by the Scandinavia-Japan Sasakawa Foundation, the Japan Eye Bank Association, the Astellas Foundation for Research on Metabolic Disorders, and the Japan Society for the Promotion of Science Overseas Research Fellowships. MT was supported by Joe, Pentti and Tor Memorial Foundation, Graduate School of Åbo Akademi University. VR was supported by Foundation of Åbo Akademi University and Magnus Ehrnrooth Foundation.

## Contributions

SV, SK and JK conceived and coordinated the study. YM, TRB, MV, MSJ, TT, SK and JK supervised the work in each contributing laboratory. Every author participated in either planning or conducting respective experiments and analysing or interpreting the data. SV, MY, VR, TA, MT, MSJ, JK wrote the manuscript. All authors approved the final version of the manuscript.

## Competing interests

The authors declare no competing interests.

## Supplementary information

### This manuscript contains Supplementary Information

**Supplementary Information 1. Supplementary file showing expression levels and statistical results of the differential expression in qualified siCTRL and siDUX4 blastomeres.** The descriptions of the columns are available at: https://github.com/shka/STRTprep/blob/v3dev/doc/result.md#outbygenediffexpxls

**Supplementary Information 2. Supplementary file showing expression levels and statistical results of the differential expression on qualified DUX4 TetOn hESC samples.** The descriptions of the columns are available at: https://github.com/shka/STRTprep/blob/v3dev/doc/result.md#outbygenediffexpxls

**Supplementary Information 3.** Supplementary table showing significantly differentially expressed enhancers in Dox (+) versus Dox (-) DUX4 TetOn hESCs.

**Supplementary Information 4.** Supplementary table showing significantly differentially expressed promoters in Dox (+) versus Dox (-) DUX4 TetOn hESCs.

**Supplementary Information 5.** Supplementary table showing DUX family proteins in primates.

**Supplementary information 6.** Supplementary table showing coordinates (KIX-9aaTAD-KBM.pdb) for modeled KIX in complex with DUX4 9aaTAD and KBM peptides.

**Supplementary information 7.** Supplementary table showing coordinates (DUX4_HD1-HD2.pdb) of DUX4 HD1-HD2 *without* bound DNA at the end of a 100 ns molecular dynamics simulation.

**Supplementary information 8.** Two concatenated supplementary movies. First, 360° view of last sampled conformation of DNA-free DUX4 (blue) from the 100 ns simulation superposed on the DNA-bound DUX4 crystal structure (red and grey) (HD1-HD2-comparison.mp4). Second, molecular dynamics simulation (100 ns) of DUX4 HD1-HD2 *without* bound DNA (HD1-HD2.mp4).

**Supplementary Information 9.** Supplementary table showing transient (BioID) and stable (AP-MS) DUX4 protein – protein interactions as well as interactions found by both BioID and AP-MS methods (both).

---

**Extended Data Figure 1.**
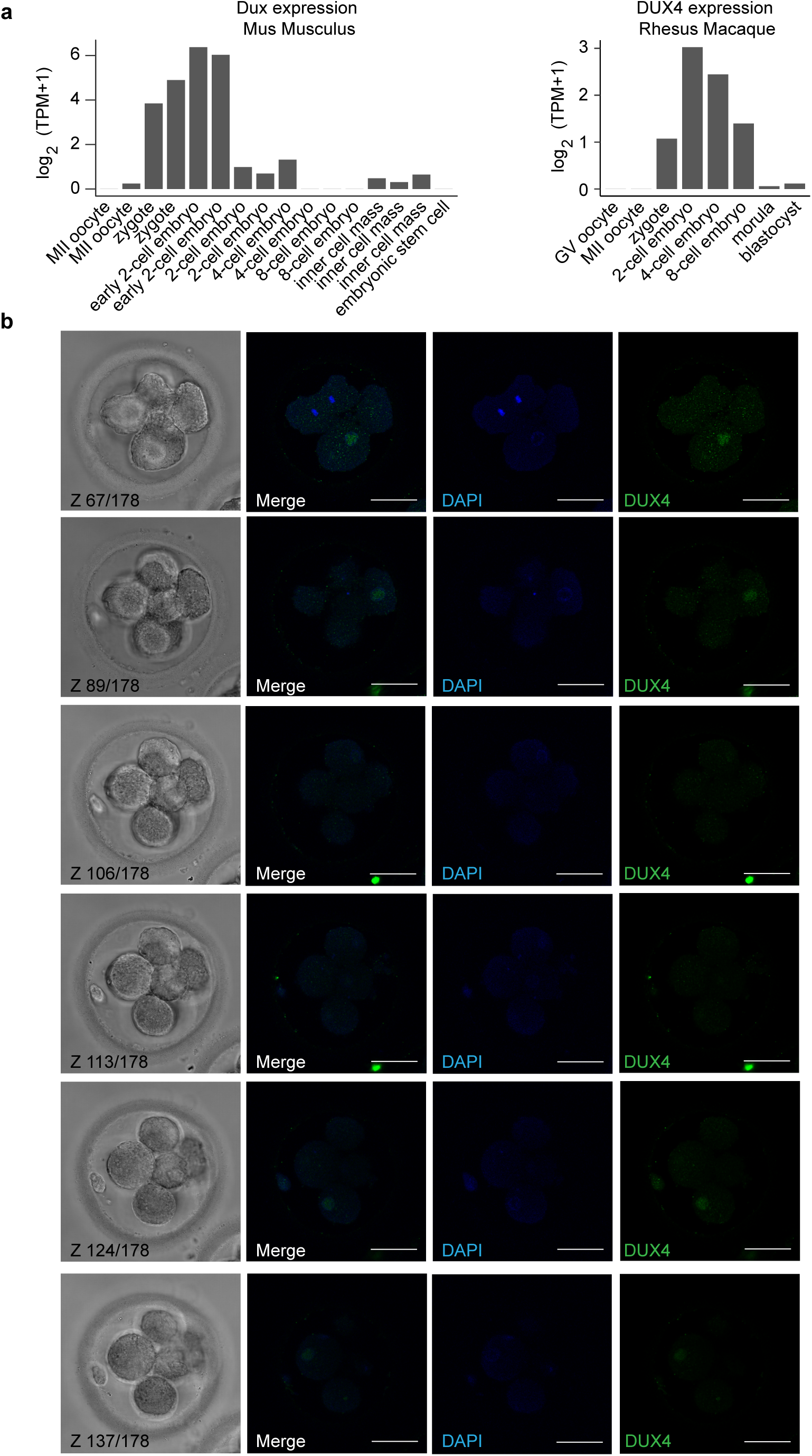
Conserved Dux/DUX4 expression in cleavage embryos. (a) Dux/DUX4 mRNA upregulation is conserved between mouse (on the left), Rhesus Macaque (on the right), and human (Fig.1b) zygotes. (b) Immunostaining of DUX4 (green) in an early 8-cell human embryo (n=1) shows heterogeneous DUX4 protein expression, consistent with observation of DUX4 clearance during the 8-cell stage. Nuclei counterstained with DAPI (blue). Six representative Z planes are shown.

**Extended Data Figure 2.**
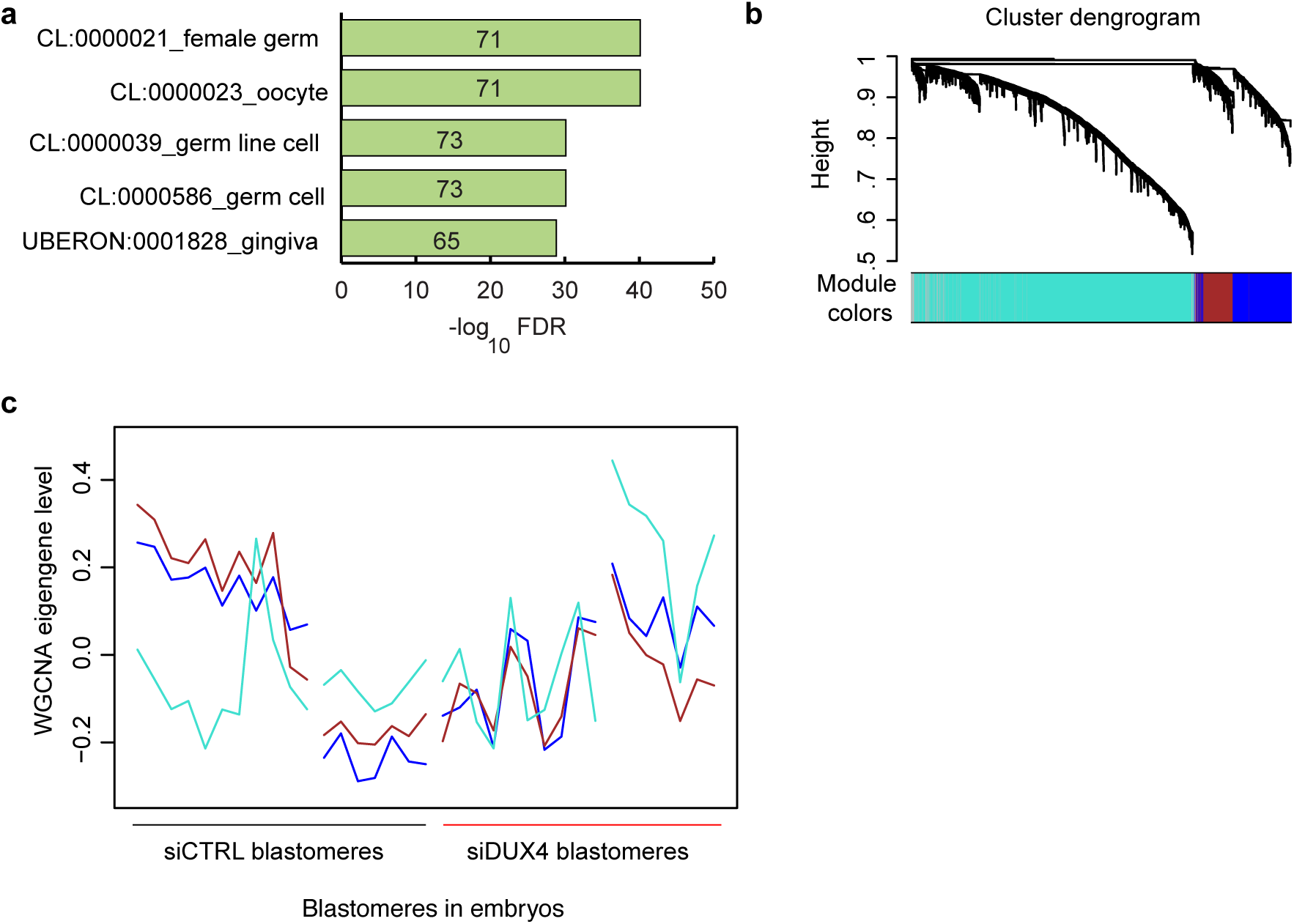
DUX4 knockdown leads to insufficient genome activation. (a) Gene expression enrichment analysis for the genes that were retained in the siDUX4 embryos using TopAnat. (b) Hierarchical clustering and module assignment of the 3,196 variable TFEs between the siCTRL and siDUX4 embryos. A maternal (turquoise) and two embryo genome activation gene modules (blue and brown) were assigned. (c) Representative expression levels of each module as indicated in (b) by the siCTRL and siDUX4 blastomeres. (d) TFEs and annotations by modules and locations.

**Extended Data Figure 3.**
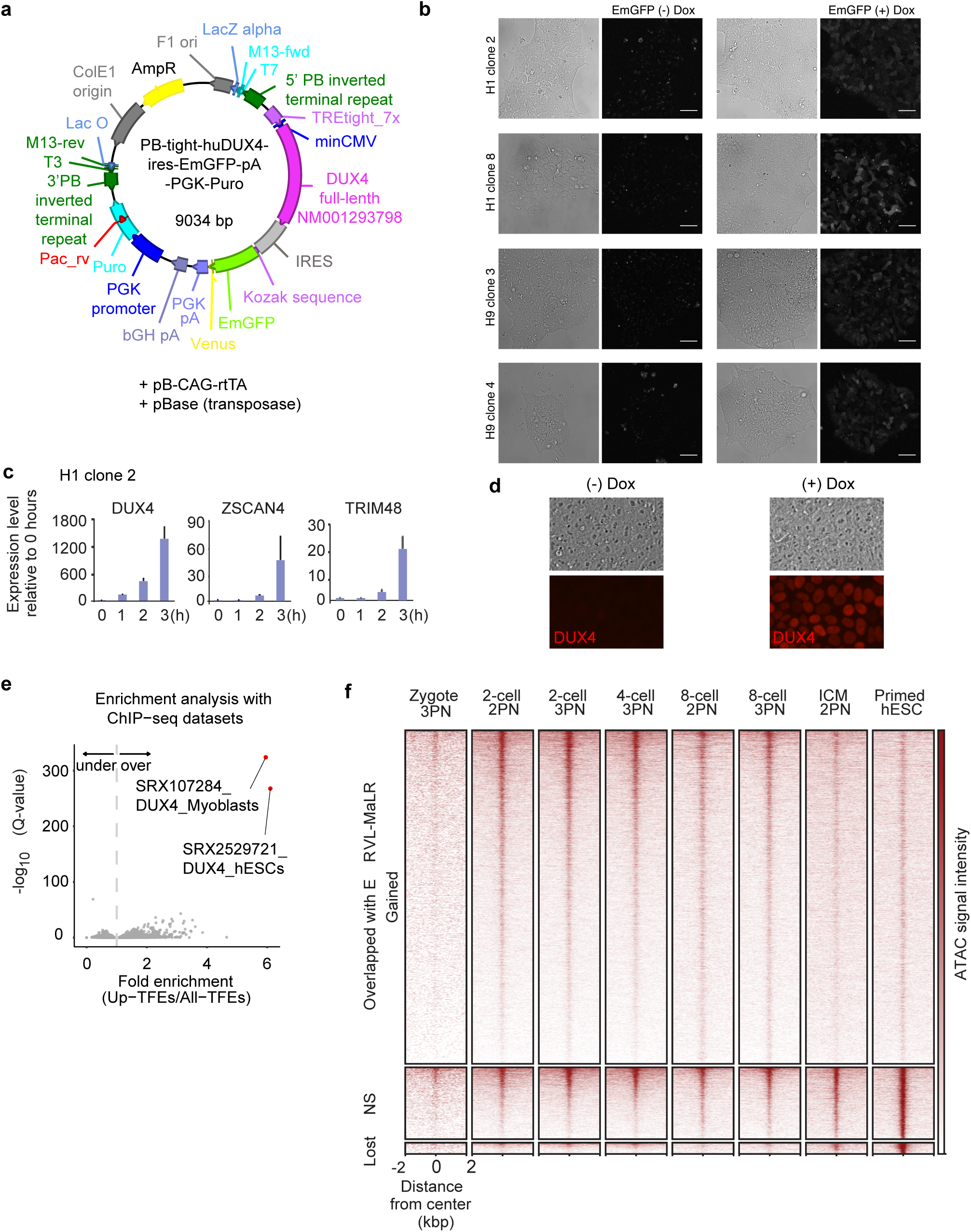
Induction of DUX4 in the DUX4 TetOn hESCs leads to expression of intergenic genome. (a) The DUX4-ires-EmGFP piggyBac vector used to establish the doxicycline inducible DUX4 TetOn hESCs in H1 (clones 2 and 8) and H9 (clones 3 and 4). (b) DUX4 TetOn hESC clones (as above) +/- 1 μg/ml doxicycline for 3 h and live imaged for EmGFP. (c) mRNA expression kinetics of DUX4, ZSCAN4, and TRIM48 after 1 h, 2 h, and 3 h doxicycline induction measured using qRT-PCR. The data for the DUX4 TetOn H1 clone 2 is shown. Similar expression patterns were also found for the H1 clone 8, and H9 clones 3 and 4. (d) DUX4 TetOn hESCs treated with 1 μg/ml doxicycline for 4 h, fixed, and immunostained for DUX4. Representative images for nuclear DUX4 staining are shown for the H1 clone 2. Similar staining pattern were seen for the H1 clone 8 and H9 clones 3 and 4. Scale bar 50 μm. (e) Enrichment analysis of the DUX4-induced TFEs with 816 publicly available ChIP-seq datasets. A total of 7,216 ChIP-seq data for transcription factors are shown. ChIP-seq data for DUX4 are shown in red. Dots on the left side of the dashed line are underrepresented, whereas dots on the right side are overrepresented. (f) ATAC-seq intensity of human early embryo around the gained, non-significant (NS), and lost ATAC-seq peaks after DUX4 induction which overlap with ERVL-MaLR elements.

**Extended Data Figure 4.**
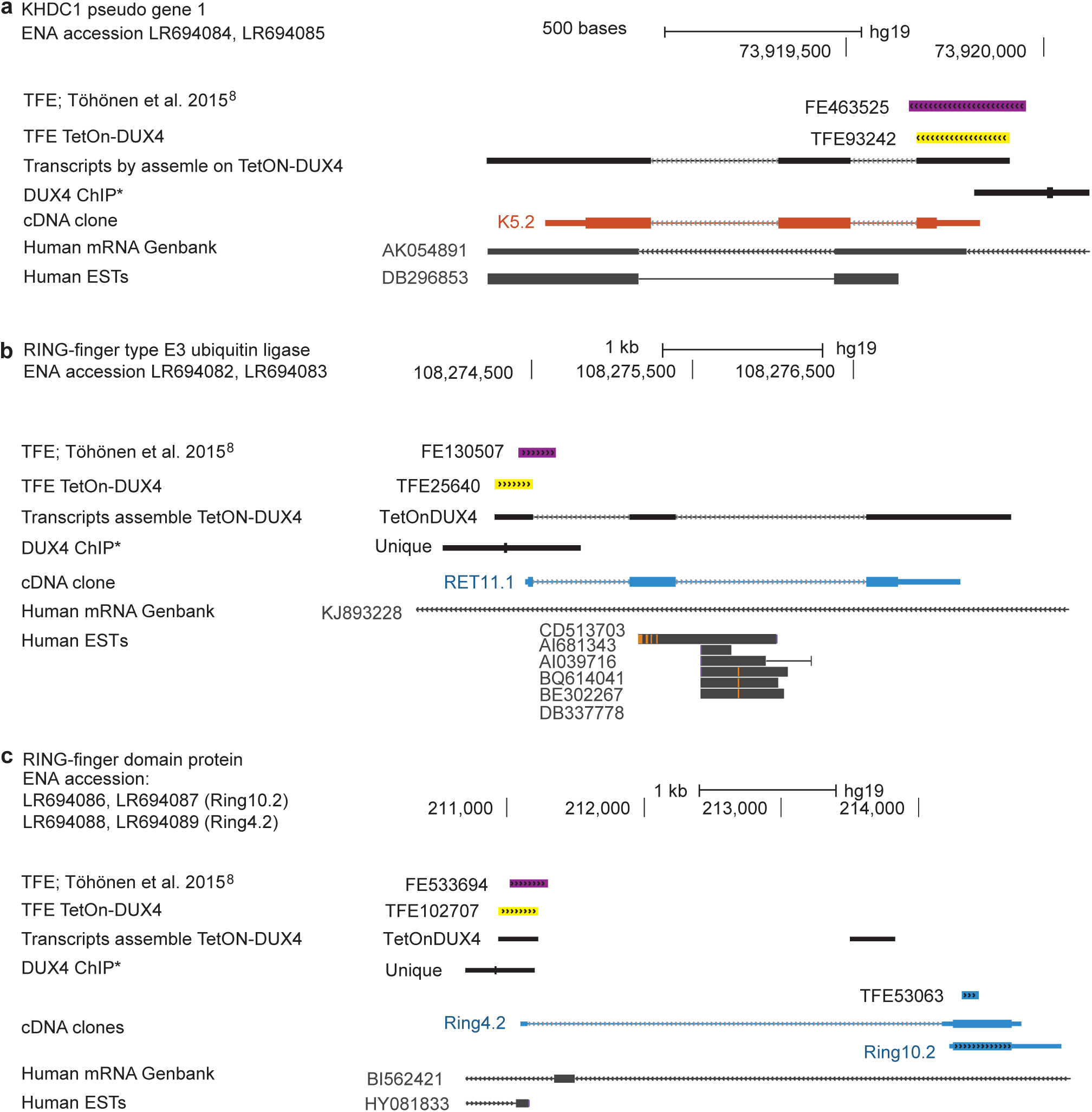
Previously unannotated putative DUX4 target genes cloned from cDNA of a human 4-cell embryo. (a) Predicted KHDC1 pseudo gene 1 (clone K5.2), at chromosome 6 (73,918,461-824 73,920,115) was expressed by human 4-cell embryos (FE463525) and upregulated in TetOn DUX4 hESCs (TFE93242). TFEs overlapped with DUX4 binding sites (DUX4 ChIP). cDNA clone K5.2 (thick orange regions indicate exons and grey thin regions indicate introns) corresponds to the KHDC1 pseudogene 1 transcript assembly in TetOn DUX4 cells. Transcript assemblies (mRNA Genbank and human ESTs), including unspliced, are shown. (b) Putative RING-finger type E3 ubiquitin ligase at chromosome 2 (108,273,771-831 108,277,850) was expressed by human 4-cell embryos (FE130507) and it was upregulated in TetOn-DUX4 hESCs (TFE25640). The DUX4 ChIP-seq peak overlapped with the TFEs. RET11.1 was cloned from human 4-cell embryo (clone RET11.1). Thick blue regions indicate exons and thin grey regions indicate introns. Transcript assemblies (mRNA Genbank and human ESTs), including unspliced, are shown. (c) Putative RING-finger domain protein at chromosome 8 (210,701-215,100) was expressed by human 4-cell embryos (TFE533694) and induced in TetOn-DUX4 hESCs (TFE102707). ChIP-seq overlapped with the TFEs. Two cDNA clones, Ring 4.2 and Ring 10.22, were expressed in the human 4-cell embyos. Thick blue regions indicate exons and grey thin regions indicate introns. Transcript assemblies (mRNA Genbank and human ESTs), including unspliced, are shown.

**Extended Data Figure 5.**
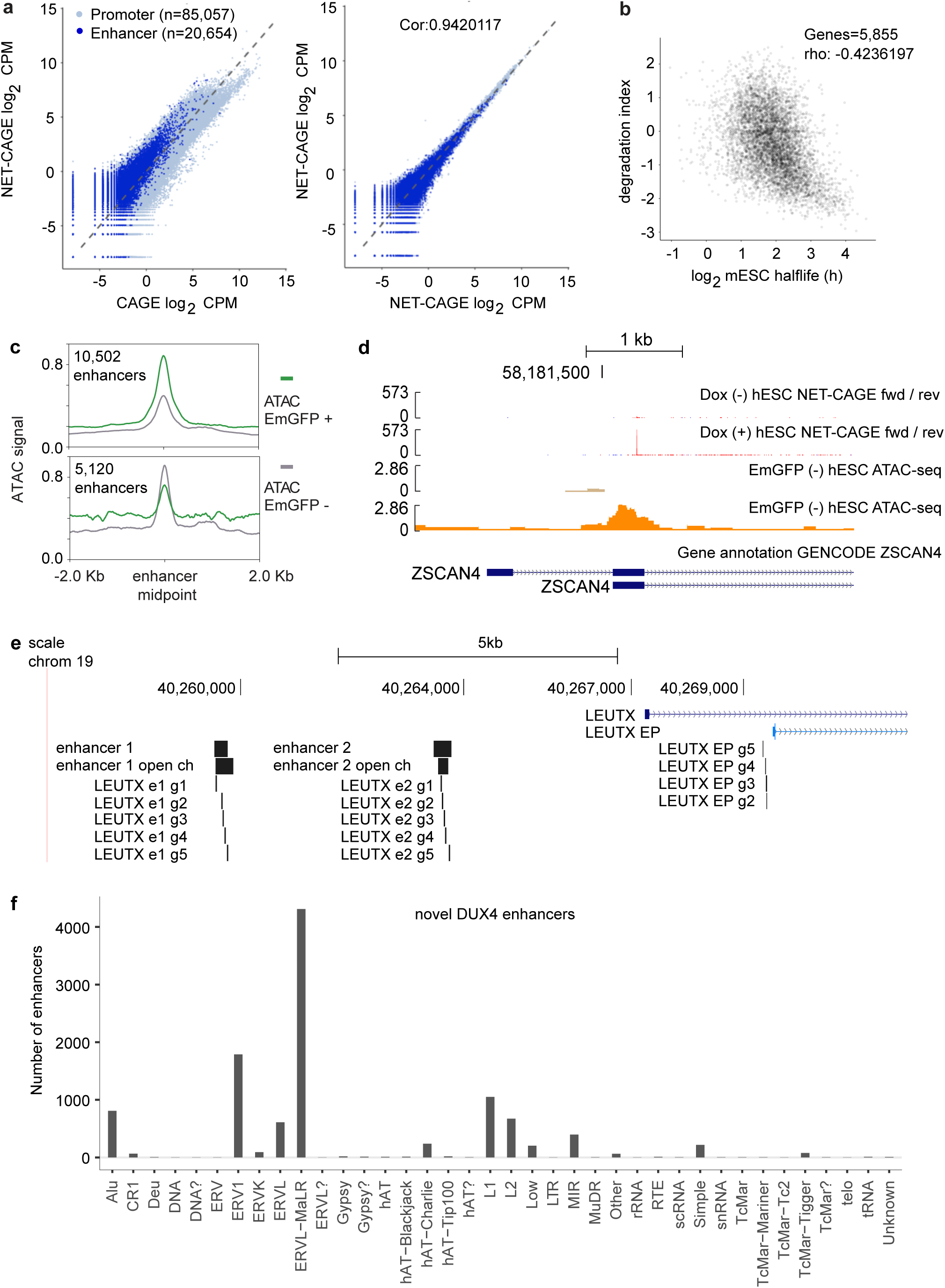
DUX4 activates novel enhancers. (a) Comparison of CAGE and NET-CAGE data in DUX4 TetOn hESCs. Unstable nascent RNA transcripts, like enhancer RNAs, were detected with high sensitivity using NET-CAGE in comparison to CAGE. (b) Scatter plot of log2 (half-lives) and degradation indices calculated as log2 (NET-CAGE/CAGE) ratios in TetOn DUX4 hESC control sample and mouse embryonic stem cells. Each dot represents a gene. (c) Metagene plots showing ATAC-seq signal for the novel DUX4 enhancers and control only enhancers in the EmGFP (+) and EmGFP (-) cells. (d) UCSC browser view of the ZSCAN4 promoter identified using NET-CAGE. NET-CAGE signal shown in the DUX4 TetOn hESCs with and without Doxicycline treatment. Chromatin status (ATAC-seq signal) shown in the EmGFP (-) and EmGFP (+) cells. (e) Positions of the CRISPR guide RNAs used for LEUTX activation. e1, enhancer 1; e2, enhancer 2; g, guide; EP, embryo promoter; open ch, open chromatin. (f) Novel DUX4 enhancers overlapping with retroelement families. All families overlapping with at least one enhancer are shown.

**Extended Data Figure 6.**
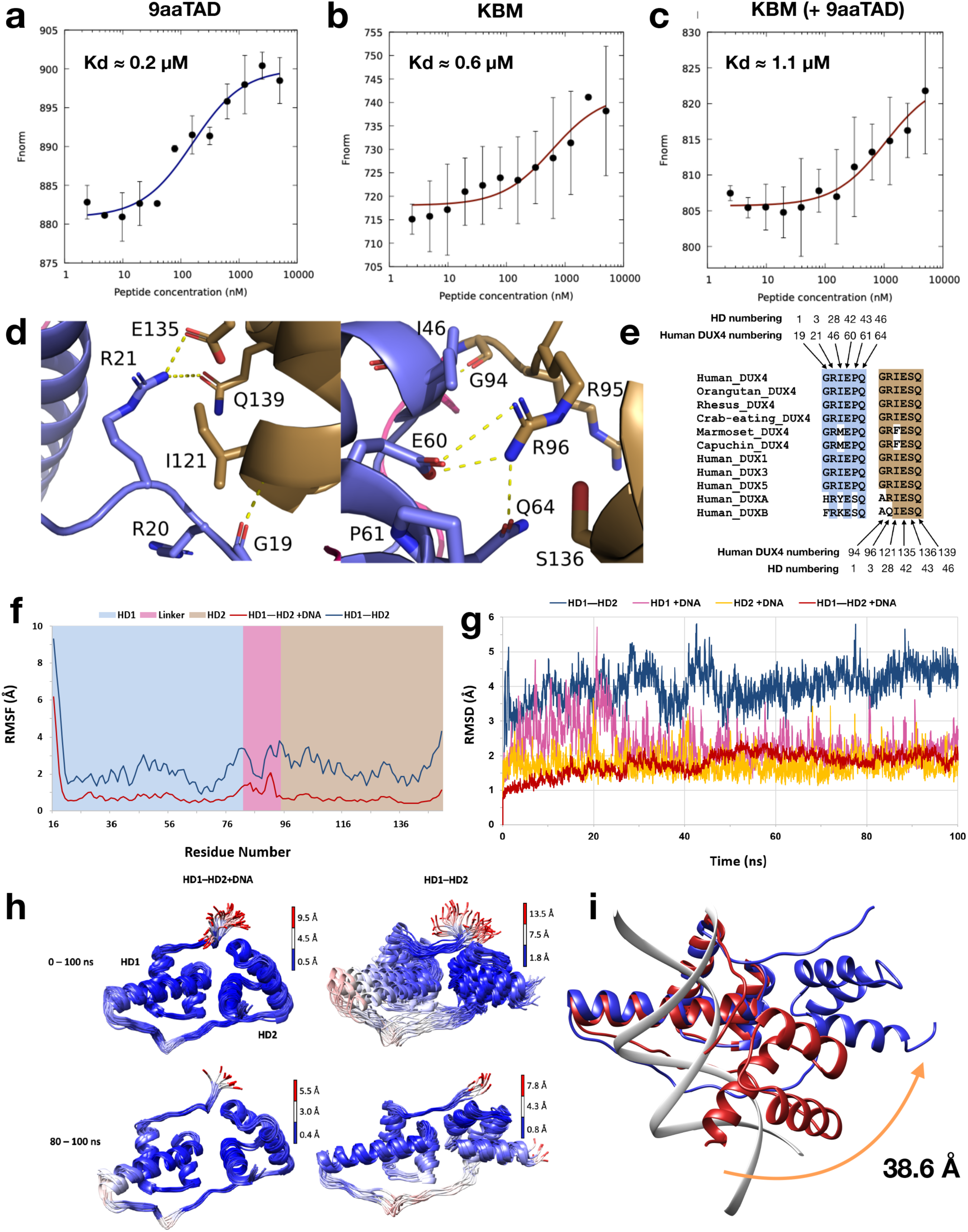
Interactions of DUX4. Microscale thermophoresis binding analysis of peptides to human KIX domain. (a) 9aaTAD peptide (C370-Q386) (b) KBM peptide (E414-E423) (c) KBM binding to KIX with saturating 9aaTAD. (d) Inter-HD interactions stabilizing DUX4 HD1 and HD2 in absence of bound DNA (e) Sequence comparison of HD1-HD2 interacting residues seen in human DUX4 with other primates and other human double HD transcription factors. (f) RMSF (Cα atoms) of X-ray structure of DUX4 with (red curve) and without (blue curve) bound DNA during a 100 ns MD simulation. HD1 (blue), linker (magenta) and HD2 (gold). (g) RMSD (backbone atoms) with reference to starting conformation) of X-ray structure of DUX4 HD1-HD2 with and without bound DNA, and separately for HD1 and for HD2 with bound DNA, during 100 ns MD simulations. (h) Superposed conformations of DUX4 with (left) and without (right) DNA, sampled during 100 ns (top) and final 20 ns (bottom) of the simulation. Chain traces are colored based on the Cα-atom RMSD relative to the median structure at 50 ns or 90 ns. DNA-bound DUX4 shows higher stability than DNA-free DUX4; both exhibit larger fluctuations at the unconstrained N-termini and linker loops. A more stable conformation of DNA-free DUX4 exposing residues of the recognition helices was attained during the last 20 ns. (i) Final pose, DNA-free DUX4 (blue), after 100 ns simulation with HD1 superposed on HD1 of DNA-bound DUX4 X-ray structure (red and grey), revealing the degree of “opening” seen in the simulation; e.g. the Cα-atom of R146 of the third helix of HD2 differs in relative position by 38.6 Å.

**Extended Data Table 7.**
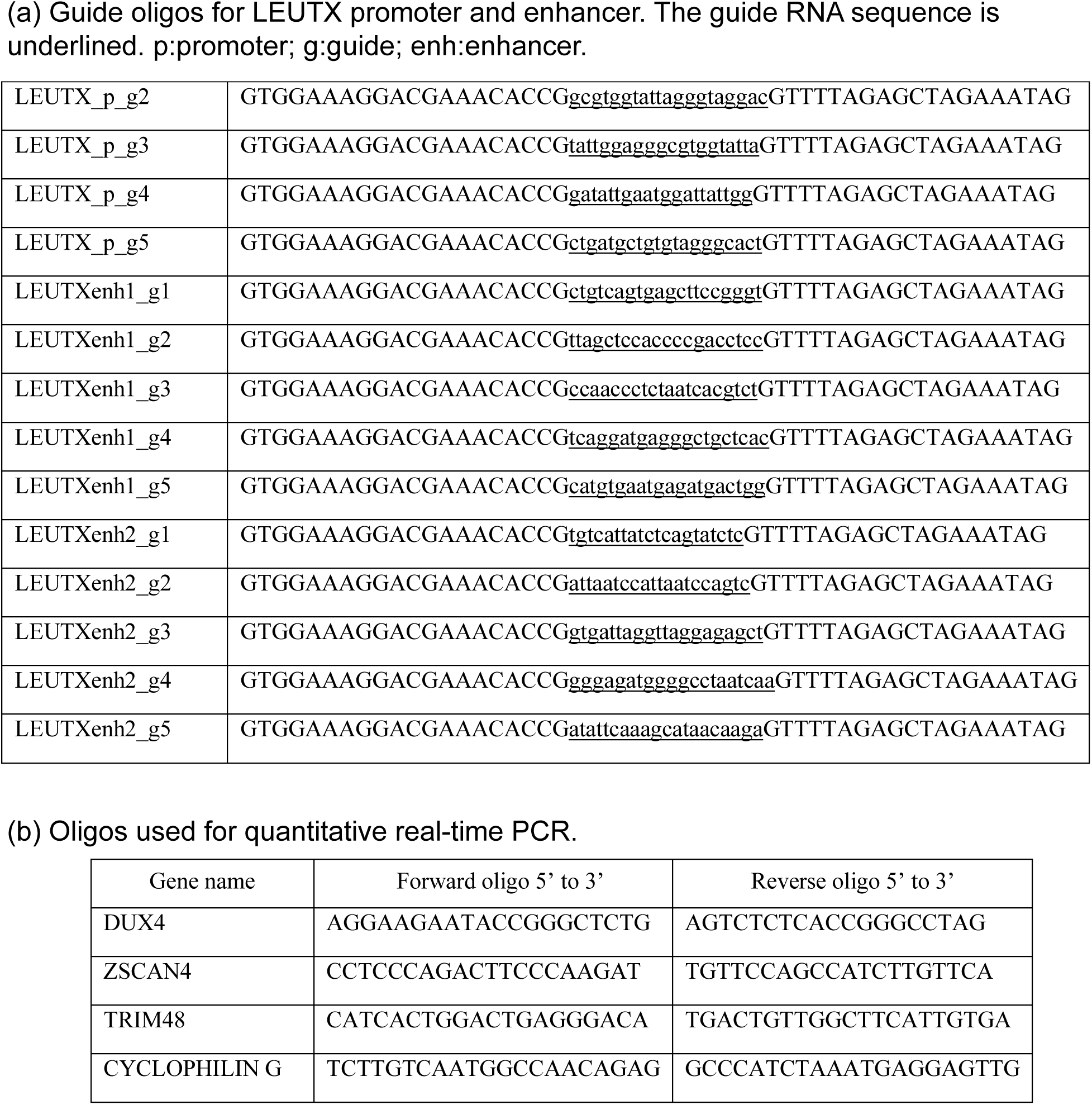
Supplementary table showing oligos that were used to (a) target LEUTX in a CRISPRa assay (b) measure expression levels of indicated genes in quantitative real-time PCR.

